# A Context-Sensitive Neural Hierarchy for Evaluating Temporal Structure in Primate Vocalizations

**DOI:** 10.1101/2025.11.14.688454

**Authors:** Ding Cui, Margaret Loewith, Audrey Dureux, Alessandro Zanini, Stefan Everling

## Abstract

Understanding how the brain encodes temporal order in communication is central to explaining how complex interactions are perceived as coherent events. In humans, disrupting the sequence of words or scenes abolishes characteristic activity in higher-order networks, but whether similar mechanisms exist in nonhuman primates remains unknown. Here we used ultra–high-field fMRI (9.4 T) in awake marmosets to test how the marmoset brain evaluates temporal structure in natural conspecific vocalizations. Animals heard vocal sequences from three social contexts (angry, conversational, food-related) presented in intact, reversed, or randomized order, with call identity held constant. Disrupting sequence order altered responses across a distributed cortical–subcortical network. Contrast to reversed order, intact sequences drove stronger activation in prefrontal, cingulate, parietal, and somatosensory regions, whereas randomization produced the most widespread disruptions, additionally recruiting motor, insular, hippocampal, and thalamic territories. Uni- and multivariate analyses revealed a core network—including prefrontal area 8, cingulate areas 24/32, somatosensory cortex, and parietal Tpt—consistently sensitive to temporal coherence, with broader recruitment under severe disruption. Network-level dynamics further varied by context: conversation elicited earlier sensitivity to sequence disruptions, angry peaked later, and food built more gradually. These findings provide the first whole-brain evidence that marmosets engage hierarchically organized, context-sensitive networks to evaluate multi-agent vocal sequence structure, establishing a cross-species bridge to human narrative processing.

## Introduction

Temporal structure is a defining feature of social communication: the order of signals shape meaning, prediction, and coherence. In humans, neuroimaging studies show that scrambling narratives—by disrupting the order of words, sentences, or movie scenes—abolishes the characteristic responses observed under intact conditions, particularly in higher-order associative regions such as the default mode and frontoparietal networks^1–13^. These findings underscore the brain’s sensitivity to sequence structure and its role in constructing coherent mental representations of events unfolding over time. But how did such sequence-processing capacities evolve? For non-human primates, “narrative-like” comprehension may involve tracking the order and outcome of social interactions—such as exchanges of calls, affiliative behaviors, or conflicts—in order to build temporally structured models of group dynamics.

Primate vocal behavior provides an especially powerful model system, as it is socially embedded and characterized by temporal structure and interactional contingency. In Old World monkeys, structured call combinations convey specific meanings, and altering sequence order modulates context-appropriate responses^14–16^. In New World marmosets, antiphonal calling exemplifies rule-based turn-taking, where response timing encodes identity, context, and coordination^17–20^. Their vocal output also exhibits hierarchical organization extending beyond adjacent call pairs^21,22^. Neurophysiological studies reinforce these behavioral findings: auditory and frontal cortices encode sequence order and hierarchical regularities, with oscillatory coupling tracking structural dependencies^23–26^; subcortical–prefrontal networks further mediate hierarchical prediction and prediction-error processing of auditory sequences^27,28^; and computational models suggest that turn-taking can emerge from temporal integration and inhibition-based control in prefrontal circuits^17,29^.

Despite converging behavioral, neurophysiological, and computational evidence for sequence sensitivity in non-human primates, no prior study has directly examined how their brains represent multi-agent vocal sequences as coherent social events, or how they respond when that temporal structure is disrupted. In contrast to the robust disruption of high-level integrative networks observed in humans processing scrambled narratives, it remains unknown whether non-human primates show similar network-level sensitivity to temporal coherence.

Here, we address this gap using ultra–high-field whole-brain fMRI in awake common marmosets (*Callithrix jacch*us) listening to natural conspecific vocalizations drawn from three distinct social contexts—angry, conversational, and food-related. Sequences were presented in intact, temporally reversed, or randomized order, while the set of individual calls was held constant. This design allowed us to isolate the contribution of temporal structure from vocal content. By combining univariate and multivariate analyses, we demonstrate that marmoset brains are broadly sensitive to vocal sequence structure, with distributed cortical and subcortical networks showing both convergent and context-dependent responses to disruption. This provides the first whole-brain evidence of sequence-based sensitivity in a non-human primate and establishes a cross-species framework for probing the neural bases of temporal coherence in social communication.

## RESULTS

### Whole-brain voxel-wise sensitivity to vocal sequence structure

During fMRI recordings in a visually dark environment, monkeys passively listened to nine conspecific vocalization stimuli (Figure 1), derived from three distinct social contexts: angry, conversation, and food. Each context was presented in three sequential formats—intact, backward, and random—while preserving the same set of individual calls. Across conditions, pooled vocalizations (collapsed across formats) evoked widespread brain activation compared to silence (block-averaged responses; Supplementary Figure 1; group-level, n = 50 runs, fixed-effects analysis). These responses encompassed prefrontal and cingulate cortices, sensory and motor regions (including auditory cortex), parietal cortex, and subcortical structures, indicating broad cortical and subcortical engagement in processing auditory social signals. We therefore focused subsequent analyses on these networks.

**Figure 1.**
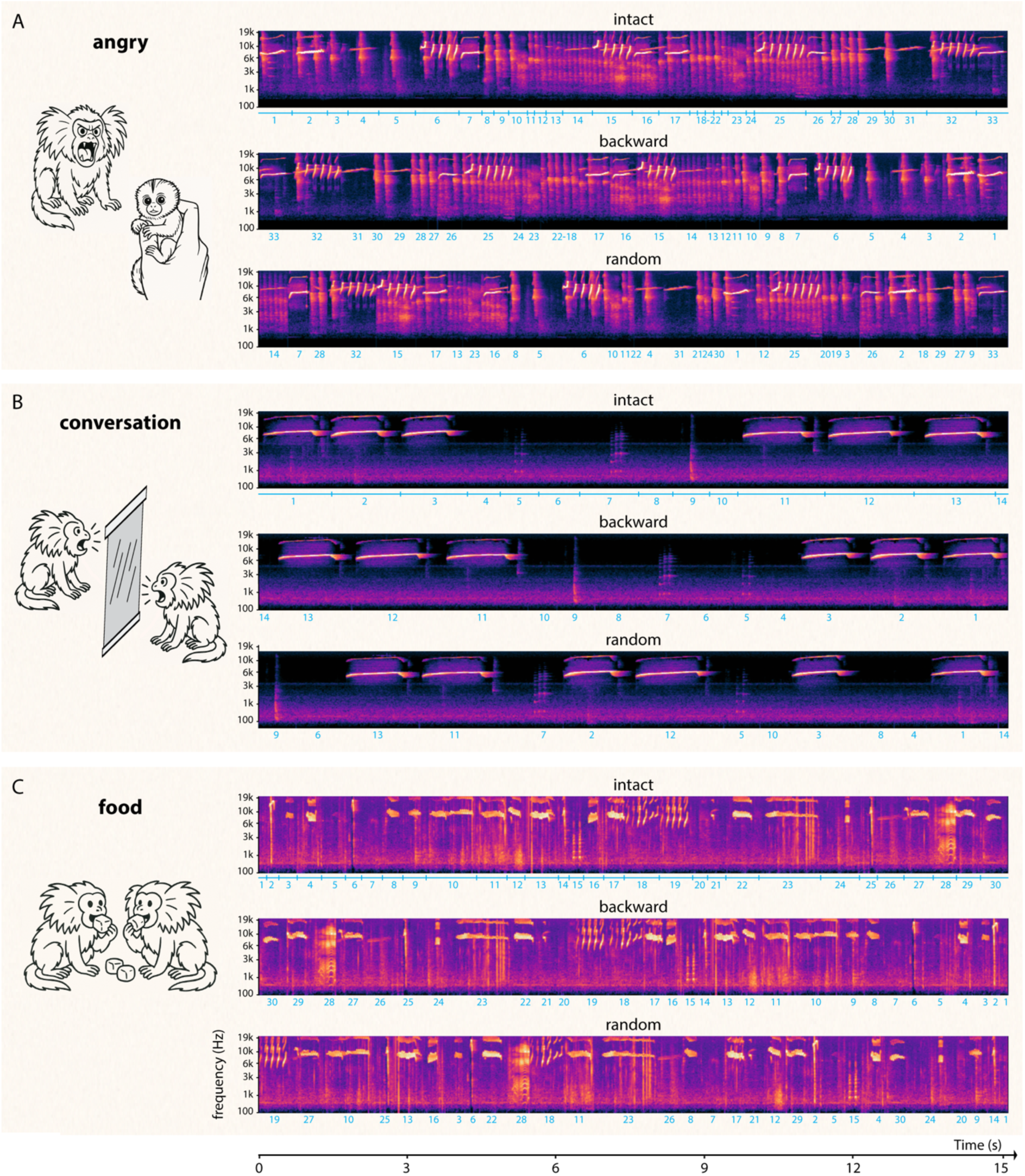
Temporal–acoustic structure of marmoset vocal sequences across social contexts. Spectrograms of the auditory stimuli used in the fMRI experiments, recorded in three behavioral contexts: (A) angry, (B) conversation, and (C) food. For each context, the same set of individual calls was presented in three sequence formats: intact (original order), backward (reversed order), and random (shuffled order). Light blue tick marks and numbers indicate segmentation into individual calls, which were reassembled to generate the scrambled versions.

To determine how the brain differentiates structured from scrambled vocalizations, we compared activations elicited by intact versus manipulated sequences (backward or random), independent of social context (Figure 2; Supplementary Figure 2). To capture unfolding vocal sequence dynamics, we modeled five successive 3s TRs within each 15s sound block using an FIR approach. Results were robust across three levels of thresholding (p<0.005; p<0.01; p<0.05; cluster size > 20); here we report findings at T > 2 (p < 0.05). Dissociations between intact and scrambled conditions emerged as early as TR1, peaked at TR2–3 (intact > scrambled), and diminished by TR4–5. Interestingly, at TR5 scrambled conditions showed slightly stronger but sparse activations than intact conditions, potentially reflecting delayed or alternative modes of processing temporally disrupted input. Spatially, the most pronounced differences occurred at TR2–3, when intact sequences induced significantly stronger activation than both backward and random sequences across bilateral cingulate regions, somatosensory cortex, and primary motor and premotor areas. Prefrontal (left-lateralized) and parietal (right-lateralized) cortices were also engaged in both contrasts, though the intact versus random comparison produced broader and more robust effects, consistent with graded sensitivity to sequence degradation, with randomization exerting the strongest disruption.

**Figure 2.**
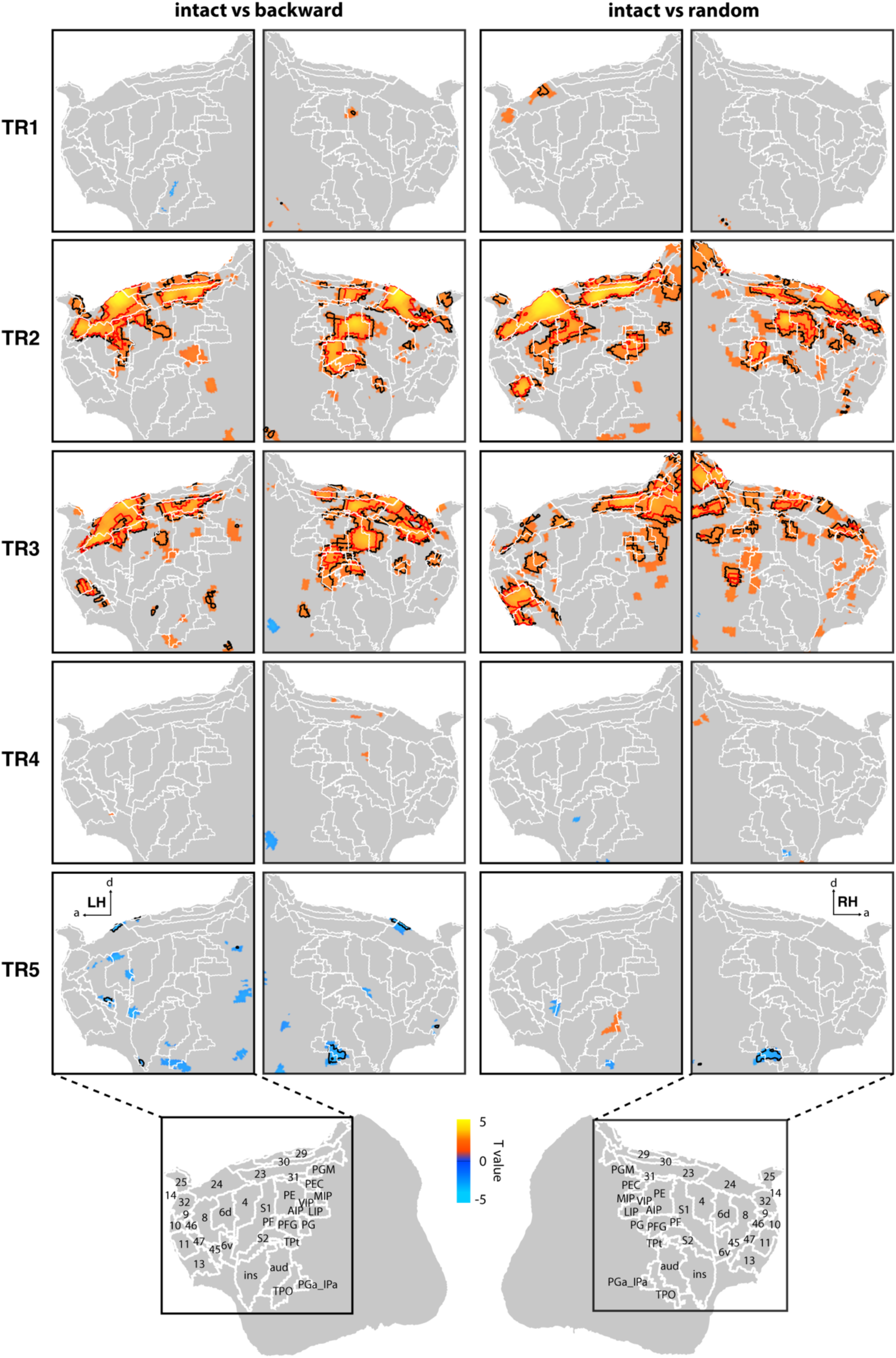
Whole-brain responses to temporally structured versus scrambled vocalizations. Group-level contrast maps (n = 50 runs, fixed effects) for intact vs backward (left) and intact vs random (right) sequences, pooled across the three behavioral contexts (angry, conversation, food). Data are shown separately for each TR (TR1–TR5) within the 15 s sound blocks. Maps are displayed on flattened, cropped hemispheric surfaces to highlight relevant cortical regions (see Supplementary Figure 2 for full-brain views). Color-coded T-maps represent surface-projected contrasts, with spatial thresholds applied from volume-based statistics: regions without outlines indicate T > 2 (p < 0.05), black outlines T > 2.3 (p < 0.01), and red outlines T > 2.68 (p < 0.005). Insets show anatomical labels from the Paxinos atlas. Abbreviations: a, anterior; d, dorsal; LH, left hemisphere; RH, right hemisphere.

### ROI-level univariate responses to vocal sequence structure in different social contexts

To refine the time-resolved whole-brain results and assess the contributions of specific regions, we conducted ROI-based analyses of univariate activation differences between intact and scrambled sequences, separately for each social context (angry, conversation, and food) at each TR (TR1-5), across a set of anatomically defined ROIs (Figure 3; Supplementary Tables 1–3). A substantial number of ROIs exhibited condition-related differences at TR2–4, typically showing stronger responses to intact than scrambled sequences, whereas fewer differences were observed at TR1 or TR5. Although most regions favored intact stimuli, some displayed stronger activation for scrambled input, particularly random sequences, at mid time points or in specific TRs. Across contexts, significant differences were more commonly observed in the intact vs random than in the intact vs backward comparisons.

**Figure 3.**
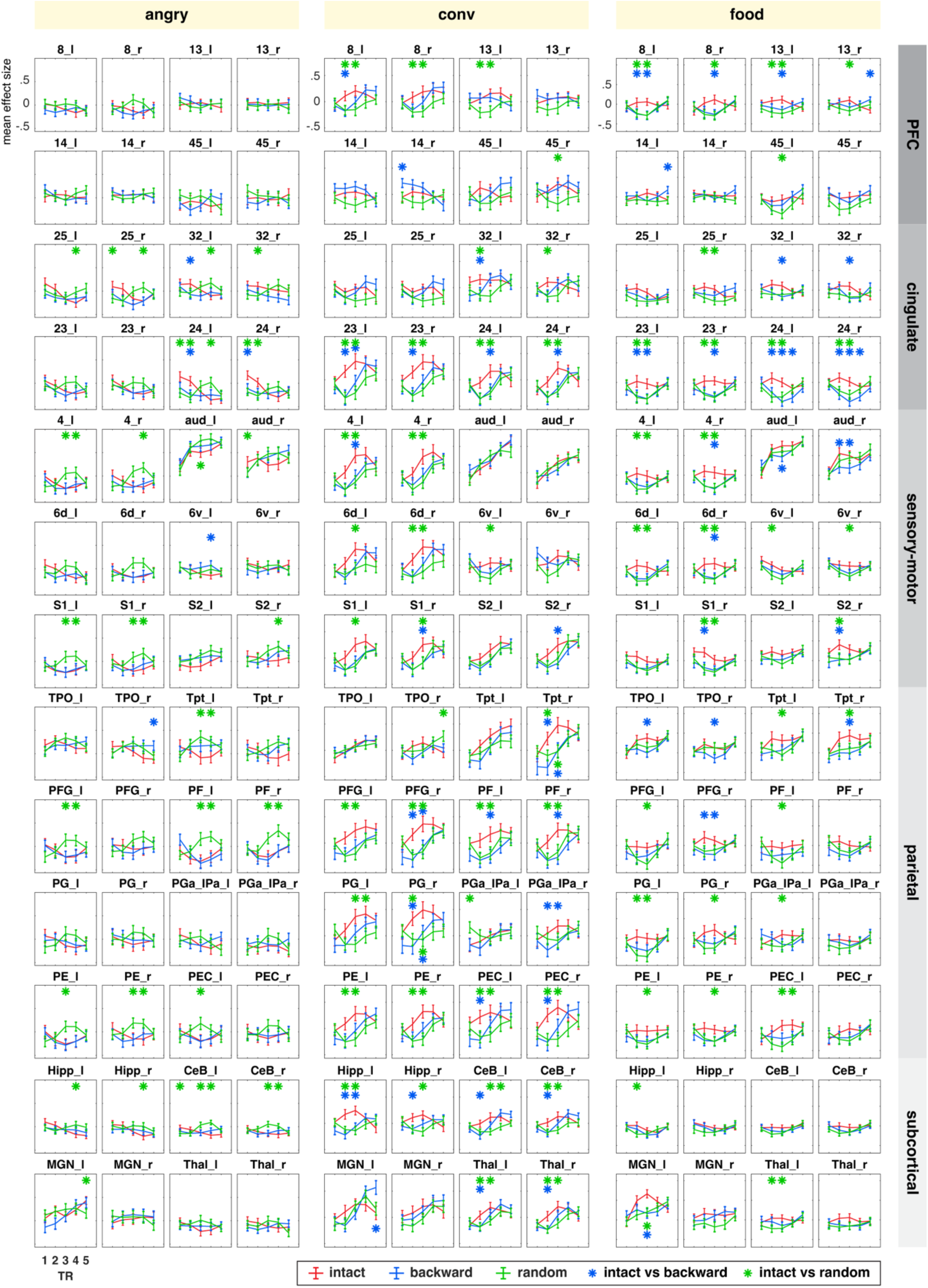
ROI-based responses to intact and scrambled vocal sequences across social contexts. Mean effect sizes (average t-values from first-level GLM coefficient maps) are shown for each ROI and TR, plotted separately for intact (red), backward (blue), and random (green) sequences in the angry, conversation, and food contexts (n = 50 runs). Error bars denote SEM. Asterisks mark significant pairwise differences based on two-tailed paired t-tests: blue, intact vs backward; green, intact vs random. ROI labels ending in “_l” and “_r” refer to left and right hemispheres. ROIs are grouped by major network (PFC, cingulate, sensory–motor, parietal, subcortical). Detailed statistics, including additional ROIs, are reported in Supplementary Tables 1–3.

The extent of these differences varied by context. In the angry context, intact versus backward differences were minimal, whereas intact versus random elicited widespread dissociations, with many ROIs showing peak activation to random sequences. In the conversation context, ROI engagement was most extensive, with numerous regions preferring intact over both backward and random sequences. The food context showed a more balanced profile, with several ROIs, including bilateral area 24, TPO, and right PFG, showing stronger effects in the intact versus backward (than versus random) contrast.

Network-wise, these results reveal distinct, context-dependent profiles. In the prefrontal cortex, structural sensitivity was most evident in the conversation and food contexts but limited in angry sequences. For example, bilateral area 8 and left-hemisphere area 13 responded more strongly to intact sequences in the conversation context, while the food context elicited more generalized intact-dominant effects. The cingulate cortex consistently differentiated structured from scrambled sequences across contexts, with bilateral area 24 showing intact preference in all three, while areas 23 and 32 varied by context. In sensorimotor regions, including area 4 and primary somatosensory cortex, bilateral engagement was observed across contexts (with the exception of left S1 in food context), while premotor area 6d showed more context-specific involvement in conversation and food context. Dissociations in these regions were particularly evident for intact versus random contrasts. The auditory cortex exhibited only subtle condition-related effects, varying slightly by context and TR. Parietal regions showed the strongest context-dependence, with nearly all parietal ROIs responsive in the conversation context, where they consistently preferred intact over scrambled stimuli. Angry and food contexts produced more restricted parietal involvement, mainly in intact versus random comparisons. Regions such as Tpt, PFG, PF, PE, and PEC were consistently recruited across all contexts, although their effect sizes varied. Finally, subcortical structures also contributed: the hippocampus reliably distinguished intact from scrambled conditions, especially random sequences, across all contexts; the cerebellum was selectively engaged in angry and conversation contexts; and the thalamus showed stronger differentiation in conversation and food context. Together, these findings indicate that neural sensitivity to vocal sequence structure is both region- and context-specific, with the conversation context eliciting the most widespread effects.

### Temporal dynamics of network-level differentiation

Based on ROI-wise univariate responses, we next quantified the proportion of significantly differentiating ROIs (out of 84) between intact and scrambled sequences at each TR, separately for each social context (Figure 4; Supplementary Table 4). In the angry context, intact versus backward comparisons showed similar engagement across TRs, while intact versus random showed low differentiation at TR1–2 followed by a sharp increase at TR3, sustained at TR4, and declining again at TR5. This suggests a delayed but transient network-level response to temporal scrambling. In the conversation context, both scrambled conditions produced a similar temporal profile, with differentiation from the intact condition peaking early at TR2–3 and reduced involvement at TR1 and TR4–5, consistent with an early sensitivity window for antiphonal vocalizations. The food context showed a more gradual build-up, with differentiation between intact and scrambled conditions increasing steadily from TR1 through TR3, peaking at TR3, and then dropping sharply at TR4–5. Despite broad similarities in temporal profiles across contexts, significant differences between intact versus backward and intact versus random emerged in all three contexts, particularly at peak TRs (angry TR3–4; conversation TR2–3; food TR3) or during the ascending phase (TR2 in food). Overall, these results reveal context-specific timescales of network engagement, with earlier peak sensitivity for conversation, later for angry, and intermediate for food.

**Figure 4.**
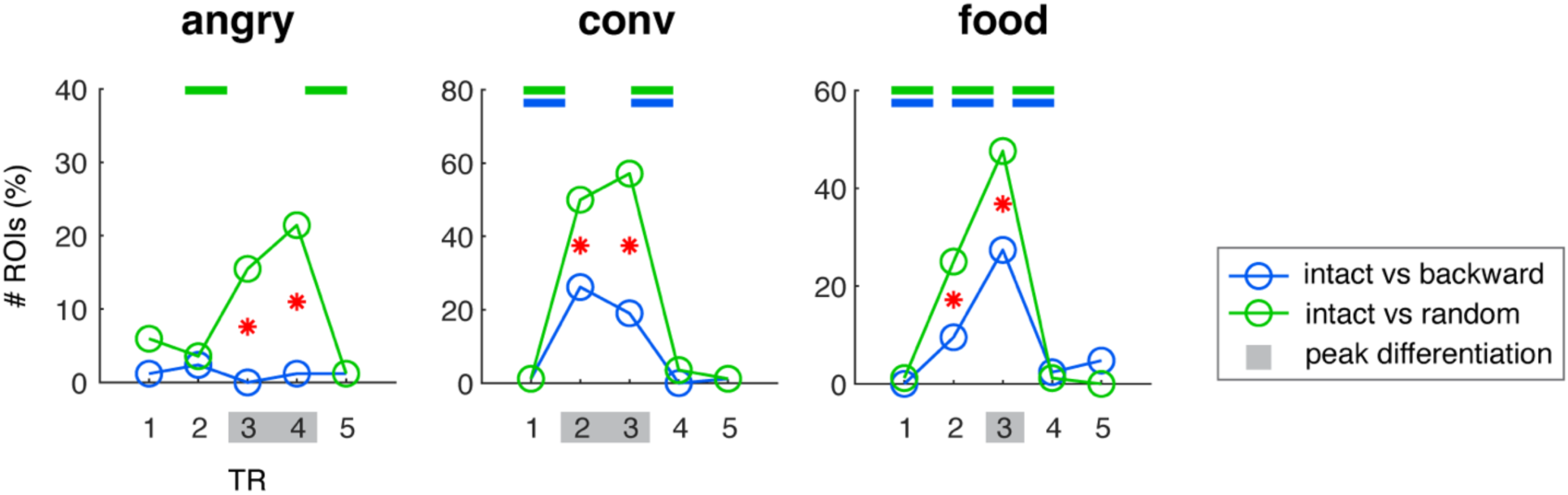
Temporal dynamics of network-level sensitivity to sequence structure. Proportion of ROIs (out of 84) showing significant differences between intact and scrambled sequences across TRs, plotted separately for angry, conversation, and food contexts. Horizontal bars above each panel mark significant changes between consecutive TRs (paired McNemar’s tests): blue for intact vs backward; green for intact vs random. Gray-shaded TRs indicate peak network differentiation. Red asterisks denote TRs where the two contrasts differed significantly from each other. Full statistical results, including all pairwise TR and contrast comparisons, are provided in Supplementary Table 4.

### ROI-level decoding of vocal sequence structure in different social contexts

To complement the univariate findings, we applied multivariate pattern analysis (MVPA) to examine representational differences between intact and scrambled conditions within all the ROIs (Figure 5; Supplementary Tables 5–7). MVPA revealed heterogeneous and distributed patterns across contrasts and contexts. Wider spreading regions exhibited significant decoding for intact versus random than for intact versus backward in the angry and conversation contexts, while the food context showed a more balanced profile. Temporally, MVPA identified significant decoding as early as TR1, especially in the angry context, and effects were distributed more evenly across TR1–5 than in the univariate analysis. Spatially, MVPA identified regions dissociating intact from scrambled conditions that were not detected with univariate analyses, including areas 8, 14, and 45 in the angry context; areas 10, 46, and 25 in conversation context; and areas 10 and 47 in food context. The insula was consistently recruited across contexts. Some regions, including areas 4, 24, 32, 30, and PFG, showed convergent effects in both univariate and multivariate analyses. Notably, intact versus random decoding consistently engaged cingulate areas 32, 24, and 30, parietal regions AIP and VIP, and the insula across all contexts, suggesting shared sensitivity to sequence disruption. Together, MVPA results extending the univariate results underscore the distributed nature of sequence sensitivity, with distinctions detectable from the earliest TRs and maintained throughout sequence presentation.

**Figure 5.**
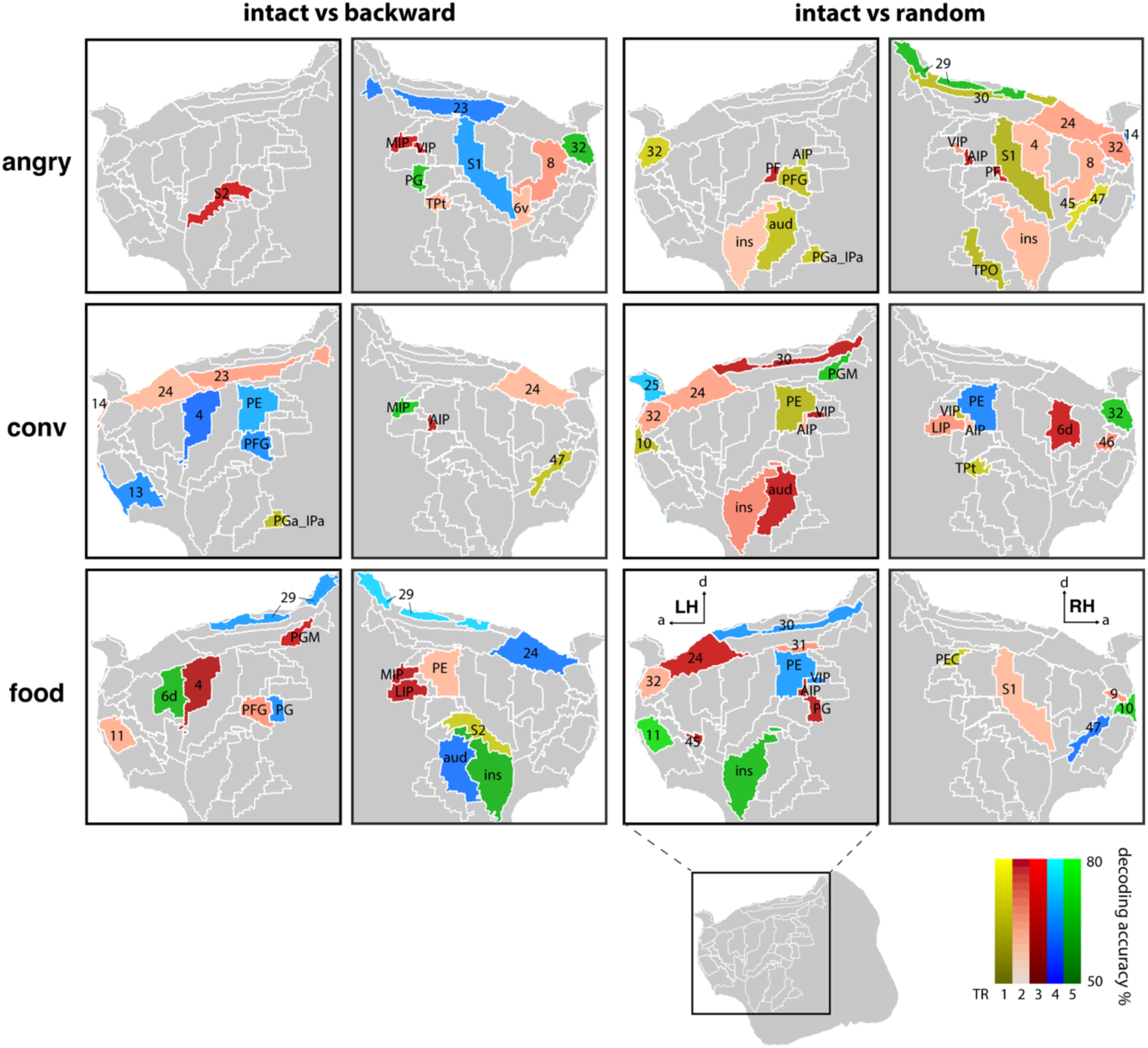
ROI-based decoding of structured versus scrambled vocal sequences. Surface maps show multivariate pattern analysis (MVPA) decoding results for intact vs backward (left) and intact vs random (right) comparisons, plotted separately for angry, conversation, and food contexts. Colors indicate TR (hue) and decoding accuracy (intensity). Only ROIs with decoding performance significantly above chance (50%) are displayed. Full decoding accuracies and statistical details are reported in Supplementary Tables 5–7.

### Distributed and convergent networks for vocal sequence processing

Taken together, univariate and multivariate results indicate that processing structured versus scrambled vocalizations engages a broad, distributed set of regions that vary with social context and degree of disruption. Figure 6A illustrates context-specific regional involvement. In the prefrontal cortex, for example, areas 9 and 11 were recruited only in food context; area 10 was engaged by intact versus random in conversation and food but not angry context; area 13 responded in conversation and food but not angry context; area 14 was recruited by intact versus backward in conversation and food, but by intact versus random in angry context; area 46 was engaged selectively in intact versus random for conversation and food context; and area 47 was involved in intact versus backward in conversation and intact versus random in angry and food context. In the cingulate cortex, areas 23, 29, and 30 were engaged by both contrasts in conversation and food context, while in angry context, area 23 responded to intact versus backward and areas 29 and 30 to intact versus random comparisons. Area 31 was absent in angry but engaged in conversation (intact vs random) and food context (intact vs backward). In premotor regions, area 6v was recruited by intact versus backward in angry context and intact versus random in conversation and food context, while area 6d was absent in angry but engaged in intact versus random in conversation and food context. In auditory cortex, intact versus random elicited responses in angry and conversation context, whereas intact versus backward effects were restricted to food context. Parietal regions also showed distinct profiles: TPO responded in all three contexts but to different contrasts, LIP was absent in angry but recruited in conversation and food context, and PGM was absent in angry but involved in conversation and food context. Subcortical regions showed context-specific recruitment as well: the amygdala responded only in angry context; the medial geniculate nucleus was engaged in angry (intact vs random), conversation (intact vs backward), and food context (both contrasts); and the cerebellum was involved in angry (intact vs random) and conversation (both contrasts), but not food context.

**Figure 6.**
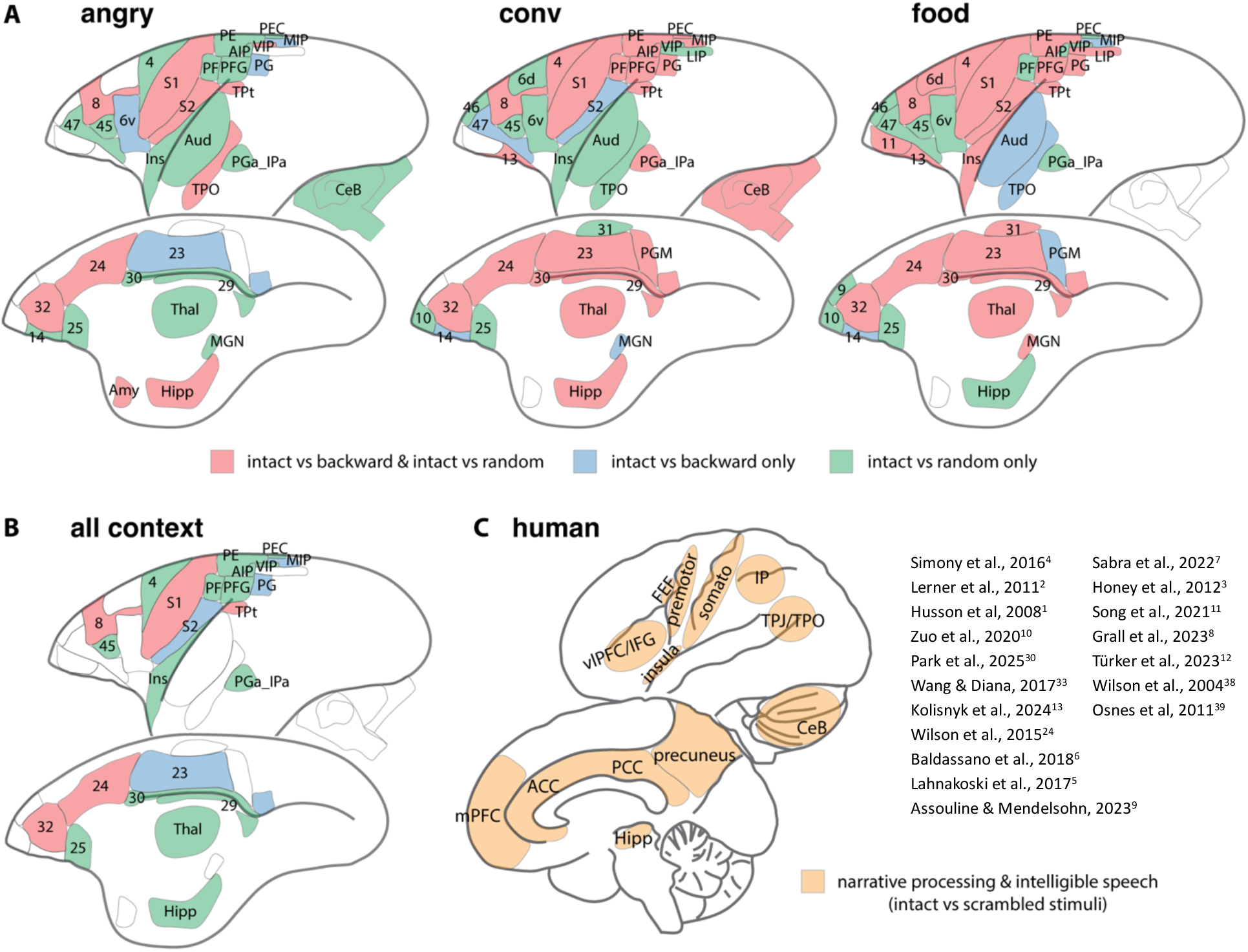
Context-specific and convergent brain networks for vocal sequence processing. Summary of ROI-level results from univariate (paired t-tests) and multivariate (MVPA) analyses comparing intact vs backward and intact vs random sequences. See Supplementary Tables 1–3 and 5–7 for full statistical results. (A) Significant ROIs plotted separately for angry, conversation, and food contexts. Blue indicates ROIs significant for intact vs backward only; green, intact vs random only; red, both contrasts. (B) ROIs consistently significant across all three contexts, using the same color scheme. (C) Reference schematic adapted from prior human studies, illustrating brain regions implicated in distinguishing structured from scrambled narrative stimuli, shown for cross-species comparison.

Despite this diversity, Figure 6B highlights a core network consistently sensitive to sequence structure across contexts, including prefrontal area 8, cingulate areas 32 and 24, primary somatosensory cortex (S1), and parietal Tpt, which responded to both contrasts. Additional convergence was observed for intact versus backward in cingulate area 23, secondary somatosensory cortex (S2), and parietal PG and MIP, whereas intact versus random recruited a broader set spanning prefrontal area 45, additional cingulate regions (25, 29, 30), motor cortex (area 4), insula, extended parietal regions (PE, PEC, PF, PFG, AIP, VIP, PGa/IPa), and subcortical structures including hippocampus and thalamus. This pattern suggests that mild temporal disruption (backward) recruits a relatively constrained network, whereas severe disruption (randomization) engages a more extensive, hierarchically organized, multimodal system encompassing sensory, associative, motor-related, and limbic regions.

## DISCUSSION

Our study provides the first direct evidence that the awake marmoset brain engages distributed cortical and subcortical networks to process the temporal structure of conspecific vocal sequences, and that this engagement is dynamically shaped by both the type of structural disruption and the social–emotional context. The organization we observe broadly mirrors systems engaged in human narrative comprehension, where temporally coherent stories or films—compared with scrambled versions—preferentially recruit prefrontal, cingulate/posterior-medial, parietal, and limbic regions beyond early sensory cortex (Figure 6C). These cross-species similarities suggest that sensitivity to temporal coherence in communicative sequences is an evolutionarily conserved feature of the primate brain that precedes the separation between New and Old World primates.

### Stabilizing sequence models and monitoring breakdown

A context-general anchor emerged in area 8, present across all contrasts (Figure 6B). In humans, dorsal frontal/FEF-adjacent cortex participates in long-timescale narrative processing, showing greater synchrony for coherent input and state shifts when comprehension fluctuates^1,4,11^. In freely moving marmosets, frontal populations exhibit sequence-sensitive, pre-stimulus state activity across conversational bouts^26^, consistent with an anticipatory control role compatible with our area8 anchor. Surrounding territories in mPFC/vmPFC support schema-like models in humans, robust for coherent input but weakened by scrambling^2,6,10^. Our results show similar context- and disruption-dependent engagement in areas 10/13/14, with area 13 particularly sensitive to intact > random in conversation and food context.

The hippocampus also plays a demand-sensitive role. In humans, vmPFC–hippocampal coupling increases when temporal order must be reconstructed^30^, and paragraph shuffling reduces default-mode-network (DMN) synchrony across listeners with hippocampal damage^10^. In macaques, medial temporal (including hippocampal) neurons integrate “what” with “when,” encoding order and recency signals^31^. Consistent with these findings, we observed hippocampal recruitment most reliably for intact > random across all contexts, and for intact > backward in angry and conversation context, suggesting that hippocampal involvement scales with ordering demands and is modulated by social–emotional context.

Lateral PFC also emerged as a key hub. Area 45 consistently differentiated intact and random sequences across contexts. Evidence from human inferior frontal gyrus (IFG)/ventral frontal operculum supports sensitivity to temporal structure encompassing modality invariance and intact-over-scrambled synchrony^13,32^, and fMRI data reveal violation exceeded consistent responses to illegal transitions^24^. Our additional findings in areas 46/47 (intact was favorable) suggest broader recruitment when temporal constraints can be exploited. Human work places lateral PFC in both regimes—long integration windows during intact narratives^3^ and heightened recruitment when order is degraded^7,33^. Although the specific effect directions vary across species and paradigms, converging evidence positions lateral PFC as a key manager of sequence structure, supporting coherence when available and promoting reassembly when it is disrupted.

The anterior cingulate sector (areas 32/24/25) was also consistently involved. In marmosets, pregenual anterior cingulate cortex (ACC; 32/24) contains a “voice patch” selective for conspecific calls and structurally coupled with auditory cortex^34,35^. Lesions to ACC reduce call variability and disrupt sequence organization^36^. Our results extend this picture: areas 32/24 were recruited across contrasts and contexts, while area 25 responded predominantly in intact > random, consistent with heightened vigilance to unpredictable input^37^. These findings resonate with human work showing dorsal ACC/pre-supplementary motor area (SMA) recruitment during comprehension breakdown^11^. Posterior cingulate and retrosplenial regions (23/29/30/31) also showed contrast-and context-dependent engagement, paralleling human PCC/precuneus sensitivity to temporal coherence^2,8^. Finally, insula recruitment for intact > random suggests a salience-switching role, as in humans where insula helps shift between DMN and attention/sensory states^4,12^.

### Routing through sensory–parietal–subcortical pathways

Unlike human A1, which is relatively invariant to scrambling^2,3^, marmoset auditory cortex distinguished intact from scrambled sequences in a context-dependent manner. This likely reflects stable low-level acoustic coding modulated by higher-order gating. Macaque neurophysiology shows similar order effects, with spiking and coupling modulated ∼450–700 ms after critical transitions^25^.

Sensorimotor recruitment further supports a prediction-monitoring account. In humans, distinction between intelligible and reversed speech engages ventral premotor and SMA, strengthening auditory–motor coupling^38,39^. In our data, premotor and somatosensory regions, and at times motor cortex, responded more to intact than scrambled sequences, despite no between-condition differences in actual vocalization (Supplementary Fig. 3), consistent with a covert sensorimotor simulation mechanism.

A key finding was robust engagement of Tpt across contexts and contrasts, with broader IPL/IPS territories recruited under randomization. This parallels macaque and human data implicating dorsal auditory–parietal pathways in vocalization selection or sequence structure, linking temporal, premotor, and prefrontal nodes^4,7,40^. In our results, Tpt acted as a cross-context hub (both contrasts), while broader IPL/IPS regions were preferentially recruited by intact > random. These patterns suggest that parietal nodes contribute additional top–down selection when sequence coherence is lost^5,11^.

Subcortical regions also modulated sequence sensitivity. The amygdala responded selectively in angry sequences, while thalamus and cerebellum showed context- and contrast-dependent involvement. These findings align with evidence that amygdala and thalamus gate social salience^34,41^, while cerebellar Crus I/II contributes to narrative timing in humans^5,9^.

### Temporal dynamics of sequence sensitivity

Beyond spatial patterns, we found distinct temporal profiles across social contexts (Figure 4). Conversation sequences elicited early peak differentiation (TR2–3), consistent with rapid integration required for antiphonal turn-taking. Angry sequences peaked later (TR3–4), suggesting longer accumulation of emotionally salient evidence. Food sequences showed a gradual build-up to TR3, then a sharp decline. Across contexts, the overall network engagement over time dissociated intact from scrambled sequences, with intact > random consistently yielding stronger and more sustained effects than intact > backward, implying that marmosets rely on both local order and global coherence with graded disruption. Together with the spatial patterns above, our findings echohierarchical temporal integration models in humans, where early sensory/motor regions track short timescales and parietal/prefrontal areas integrate over longer windows^1,3^.

Future studies combining naturalistic multi-individual recordings with fMRI and electrophysiology could map region-wise integration timescales and test whether the human lag-gradient (A1 → temporal → parietal → precuneus/mPFC) has a marmoset counterpart, and how social context reconfigures it.

## Conclusion

Taken together, our findings demonstrate that the awake marmoset brain recruits a distributed, hierarchically organized, and context-sensitive network to process vocal sequence structure. Area 8 provides a context-general control anchor; mPFC/vmPFC and hippocampus consistent with long-timescale modeling and order reconstruction; lateral PFC appears to manage coherence and reassembly; cingulate and insula are consistent with roles in valuation and state switching; a Tpt-centered dorsal auditory–parietal–premotor pathway likely maintains and updates order-sensitive predictions; and subcortical relays may contribute salience and timing. Temporal profiles vary with social context and intensify under randomization. These features suggest that mechanisms for integrating coherent sequences, monitoring breakdowns, and routing predictive information are present in a New World primate, supporting an evolutionarily conserved sensitivity to temporal coherence in communicative sequences.

## METHODS

### Subjects

Six common marmosets (*Callithrix jacchus*; 2 females, 4 males; 49–81 months) participated in the fMRI experiments. All procedures conformed to the guidelines of the Canadian Council on Animal Care and were approved by the University of Western Ontario Council on Animal Care. Each animal was implanted with a PEEK head post following Zanini et al.^42^. After a 2-week recovery, animals underwent ∼3 weeks of habituation to head-fixation and the MRI environment.

### Auditory stimuli and experimental design

Natural vocalizations were recorded from small colonies comprising several marmoset groups housed in separate rooms (2–6 animals per cage, 4–10 cages per room). With one exception, recorded animals did not participate in the fMRI study and were housed separately from scanning subjects. Recordings were made in three behavioral contexts: (1) **Angry:** an infant was briefly separated for veterinary exam; the microphone was near the infant but captured adult calls from the room; (2) **Conversation:** two non–cage-mate adults were placed in separate small cages in an experimental room; after an initial period with mutual visibility, a screen blocked visual contact; the microphone was centered between cages to capture antiphonal phee exchanges; (3) **Food:** three adults in one cage received marshmallows; the microphone was positioned in front of the cage during shared feeding.

Spectrograms (Audacity v3.6) were inspected to select one clear ∼15 s segment per context. The “angry” segment contained infant vocalizations with aggressive adult calls; “conversation” captured one turn-taking exchange of phee calls; “food” contained dense affiliative vocalizations. To attenuate low-frequency noise while preserving vocal content, we applied a high-pass filter (angry/food: 500 Hz; conversation: 300 Hz) and normalized spectral power (Audacity “Normalize,” default settings). Each ∼15 s segment was then hand-segmented into individual calls; the same calls were reassembled in reverse (backward) or randomized order, preserving call identity but altering sequence structure. Overlaps (e.g., infant phee overlapping short adult calls) were segmented according to the most salient call (e.g., infant phee). In total, nine stimuli were created: 3 contexts × 3 sequence formats (intact, backward, random).

Stimuli were presented in a block design. Each run comprised 36 blocks alternating silence and sound (15 s each), and followed by an additional silence block at the end of the run (total 555 s; 185 volumes). The nine stimuli appeared twice per run. Ten pseudorandomized block orders minimized consecutive presentations of similar stimuli and were counterbalanced across subjects and runs.

### fMRI data acquisition

Preparation and setup followed Zanini et al.^42^. Awake animals were head-fixed in a custom 3D-printed MRI-compatible chair (sphinx posture). Sounds were delivered in the dark via Sensimetrics S14 earphones secured with earplugs and bandage and controlled by custom MATLAB (R2018b) code. An MRI-compatible camera (12M-i, MRC Systems) recorded frontal face videos (Raspberry Pi 3B+, custom Python 3.9). A transistor–transistor logic (TTL) box synchronized EPI onset with audio and video.

Scanning used a 9.4 T horizontal scanner (31 cm Varian magnet; Bruker Avance NEO console) with a custom 15 cm gradient coil and an 8-channel receive coil inside a quadrature birdcage transmit coil^43^. Functional data were acquired with gradient-echo EPI: TR = 3 s, acquisition time = 1.5 s, TE = 15 ms, flip angle = 40°, field of view = 48 × 64 mm, matrix = 96 × 128, voxel size = 0.5 mm isotropic (0.5 × 0.5 × 0.5 mm), 42 axial slices, bandwidth = 400 kHz, GRAPPA = 2 (L–R). Phase-encoding was alternated within sessions (L–R and R–L). To reduce masking by scanner noise, we used a continuous acquisition with a built-in silent interval^44^: each TR included 1.5 s of silence followed by 1.5 s of whole-brain acquisition. For each animal, a T2-weighted structural image was acquired (TR = 7 s, TE = 52 ms, FoV = 51.2 × 51.2 mm, bandwidth = 50 kHz, resolution = 0.133 × 0.133 × 0.5 mm). Each session lasted ∼60 min. Subjects completed 1–3 sessions (total 50 runs: M11 = 10, M13 = 5, M15 = 6, M16 = 10, M19 = 5, M20 = 14).

### Preprocessing

Preprocessing used AFNI^45^ and FSL 6.0.7^46^. Raw DICOMs were converted to NIfTI (dcm2niix), reoriented (FSL fslswapdim, fslorient), and distortion-corrected with topup/applytopup using reverse phase-encoded EPI. Transient spikes were attenuated with AFNI 3dDespike. Slice-timing correction used 3dTshift (-tzero 0). Motion correction used 3dvolreg (realignment to the middle volume). Data were spatially smoothed with a 1.5 mm FWHM Gaussian kernel (3dmerge) and temporally band-pass filtered 0.01–0.1 Hz (3dBandpass).

### Whole-brain GLM

We estimated hemodynamic responses with AFNI 3dDeconvolve. Low-frequency trends were removed via polynomial regressors. First, to assess overall averaged activation to vocal blocks, we modeled responses with BLOCK(15,1). Second, to characterize within-block dynamics, we used a finite-impulse-response (FIR) basis TENT(0,15,6), yielding five TR-resolved estimates (every 3 s) within each 15 s sound block. Resulting T-maps were aligned to each subject’s skull-stripped T2 image (FSL FLIRT) and nonlinearly registered to the NIH marmoset template^47^ using ANTs ApplyTransforms. Group analyses were conducted with AFNI 3dttest++ using fixed effects across runs (n = 50). One-sample two-sided t-tests generated condition-versus-silence maps (Supplementary Fig. 1); paired two-sided t-tests compared conditions (Figure 2; Supplementary Figs. 2). For visualization, we show thresholds at T = 2 (p < 0.05), 2.3 (p < 0.01), and 2.68 (p < 0.005), each with a minimum cluster size of 20 voxels.

Surface visualizations were rendered on the flattened NIH template using Connectome Workbench 2.0. Because volume-to-surface mapping can attenuate statistical values^48,49^, thresholding was performed in volume space; thresholded volumes were converted to surface masks and intersected with unthresholded surface maps to preserve the original volume-based cluster topology while displaying continuous surface values. Anatomical labels followed the Paxinos atlas^50^.

### Univariate region-of-interest (ROI) analysis

Cortical and subcortical ROIs were defined on the NIH template^47^ using Paxinos-based parcellations^50^. Using the pooled-vocalization activation map as a guide (Supplementary Figure 1), we focused subsequent ROI analyses on all parcels in prefrontal, cingulate, somatosensory/motor, parietal, and key subcortical regions (including cerebellum). For adequate voxel counts per ROI—particularly for MVPA—sub-parcels were merged into 84 bilateral ROIs (Supplementary Tables).

For each run, condition, and TR, voxel-wise T-values from first-level GLMs were averaged within each ROI. Two-sided paired t-tests compared intact vs backward and intact vs random within each context (angry, conversation, food) at each TR (Figure 3; Supplementary Tables 1–3). We report p < 0.05 and provide FDR-corrected results as complementary evidence. Given the exploratory, tuning-like mapping across >80 anatomically defined ROIs and expected heterogeneity across contexts, we present uncorrected results to avoid excessive Type II error, interpreting them in concert with whole-brain maps and theory; FDR-corrected outcomes are reported alongside.

To assess network-level temporal dynamics (Figure 4; Supplementary Table 4), we converted each ROI test to a binary outcome (significant p < 0.05 vs not) and applied continuity-corrected McNemar tests. (i) Across-time: within each contrast, all TR pairs were compared to test whether the number of engaged ROIs changed over time. (ii) Across-contrast: at each TR, intact vs backward was compared to intact vs random. P-values were FDR-corrected across tests.

### Multivariate ROI-based decoding

Pairwise MVPA compared intact vs backward and intact vs random within each context and TR using The Decoding Toolbox^51^ with a linear SVM. For each ROI, GLM T-values (per run, condition, TR) served as features. We used leave-one-run-out cross-validation; mean accuracy across folds was the decoding score. Significance was assessed by 1,000-permutation label shuffling within ROI with identical cross-validation; p-values reflected the position of the observed accuracy in the null distribution. Results were considered significant at p < 0.05; FDR-corrected outcomes are provided as complementary evidence (Figure 5; Supplementary Tables 5–7).

### Analysis of mouth movement

Although acoustic confirmation of vocal output was not possible during scanning, we quantified visible mouth movements as a proxy for potential vocalizations to evaluate contributions to somatomotor activity (Supplementary Fig. 3). Videos (frontal face view within the circular head coil) were processed with DeepLabCut 2.3.8 (Python 3.8). Five representative videos (distinct monkeys with clear mouth motion) provided 25 labeled frames each (125 total) for training (100,000 iterations). Upper and lower lip landmarks were tracked; vertical distance between landmarks was computed over time. Using synchronization with fMRI TRs and block onsets/offsets, distance traces were segmented by condition and baselines. Mouth events were defined as peaks above a baseline threshold (no-movement state >90% of samples). For each condition, we counted peaks within each sound block and the immediately following silent block and expressed counts as a percentage of blocks for that condition.

## Acknowledgements

Support was provided by a Discovery grant by the Natural Sciences and Engineering Research Council of Canada and the Canadian Institutes of Health Research (FRN 183973). We also acknowledge the support of the Government of Canada’s New Frontiers in Research Fund (NFRF), [NFRF-T-2022-00051]. We wish to thank Rebekah E Gilliland for providing the recordings used as stimuli in this study, Cheryl Vander Tuin, Whitney Froese, Hannah Pettypiece, and Miranda Bellyou for animal preparation and care, Dr. Alex Li and Trevor Szekeres for scanning assistance, Dr. Kyle Gilbert, and Peter Zeman for coil designs.

## Supplementary Figures

**Supplementary Figure 1.**
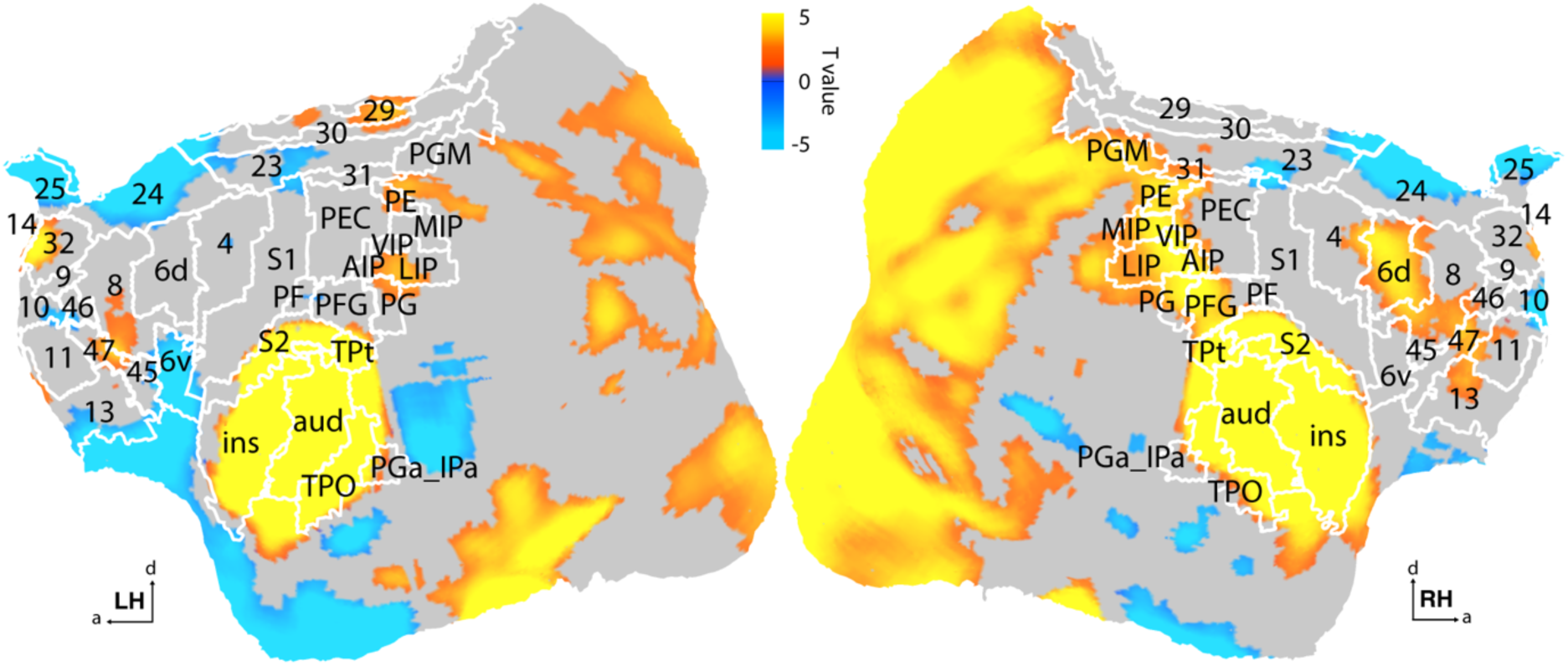
Group-level (n = 50 runs, fixed-effects) contrast maps are shown for vocalization vs baseline (silence). Color-coded T-maps reflect surface-based contrast values, with spatial thresholding applied based on corresponding volume-based statistical maps: T > 2 (p < 0.05). a - anterior, d - dorsal, LH - left hemisphere, RH - right hemisphere.

**Supplementary Figure 2.**
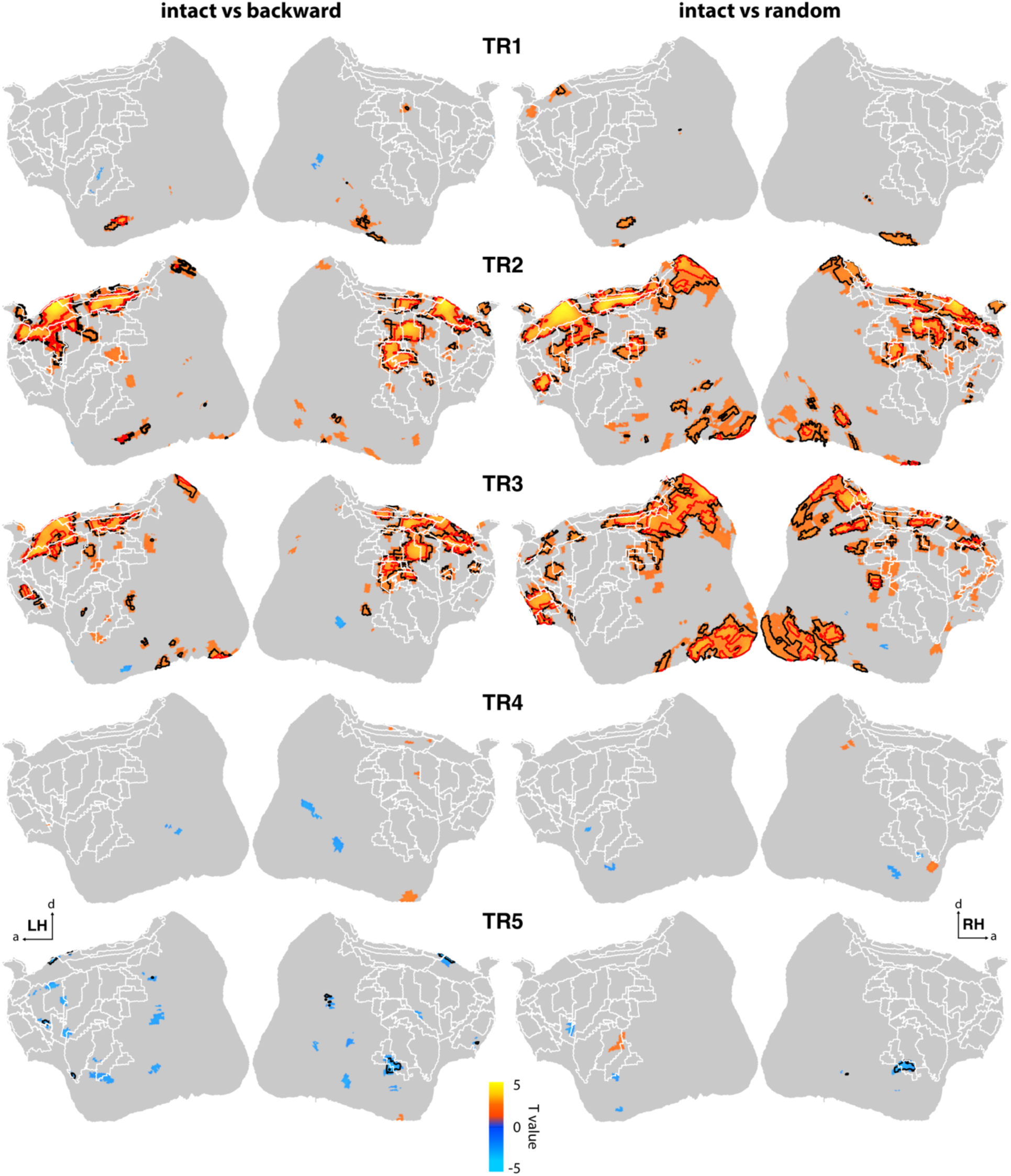
Full whole-brain maps for Figure 2.

**Supplementary Figure 3.**
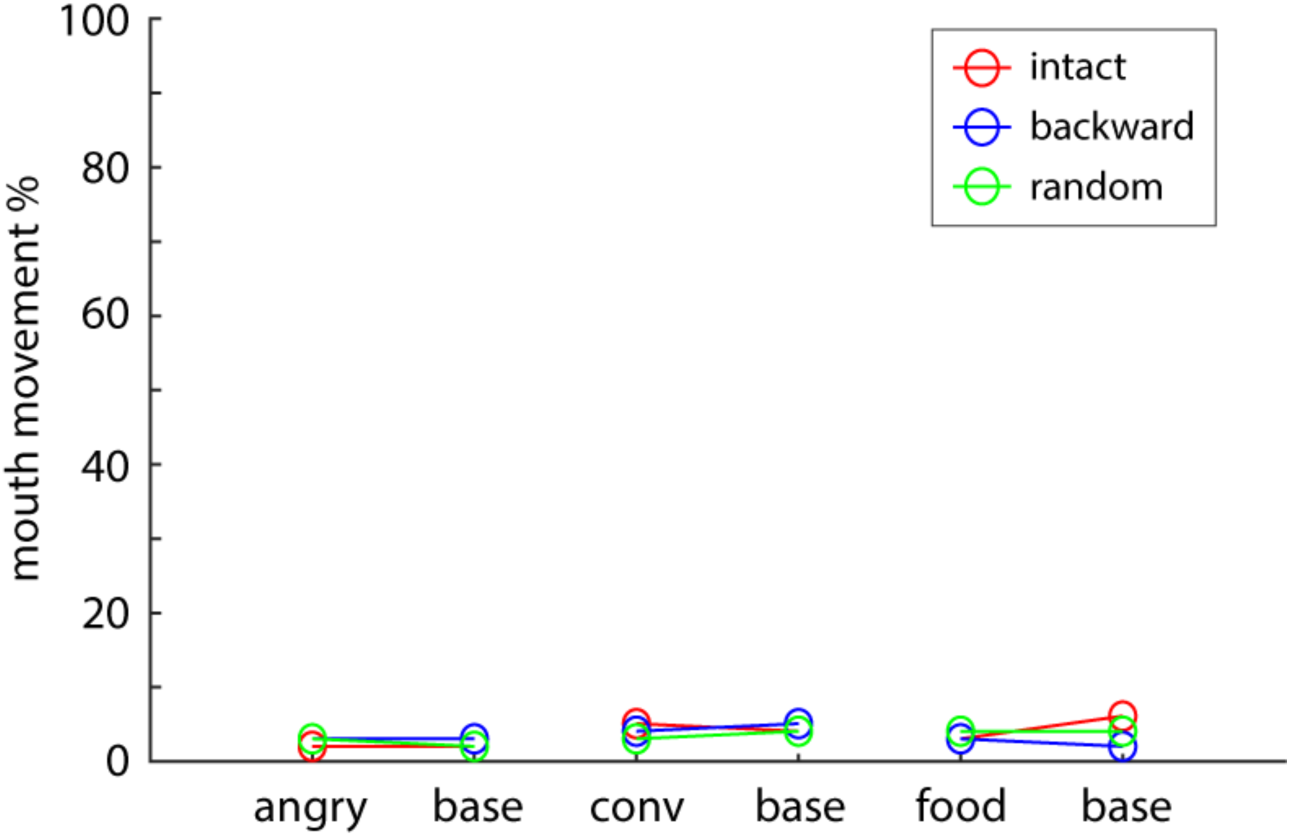
Mouth movements as potential effort for vocalizations for each condition.

**supplementary Table 1.**
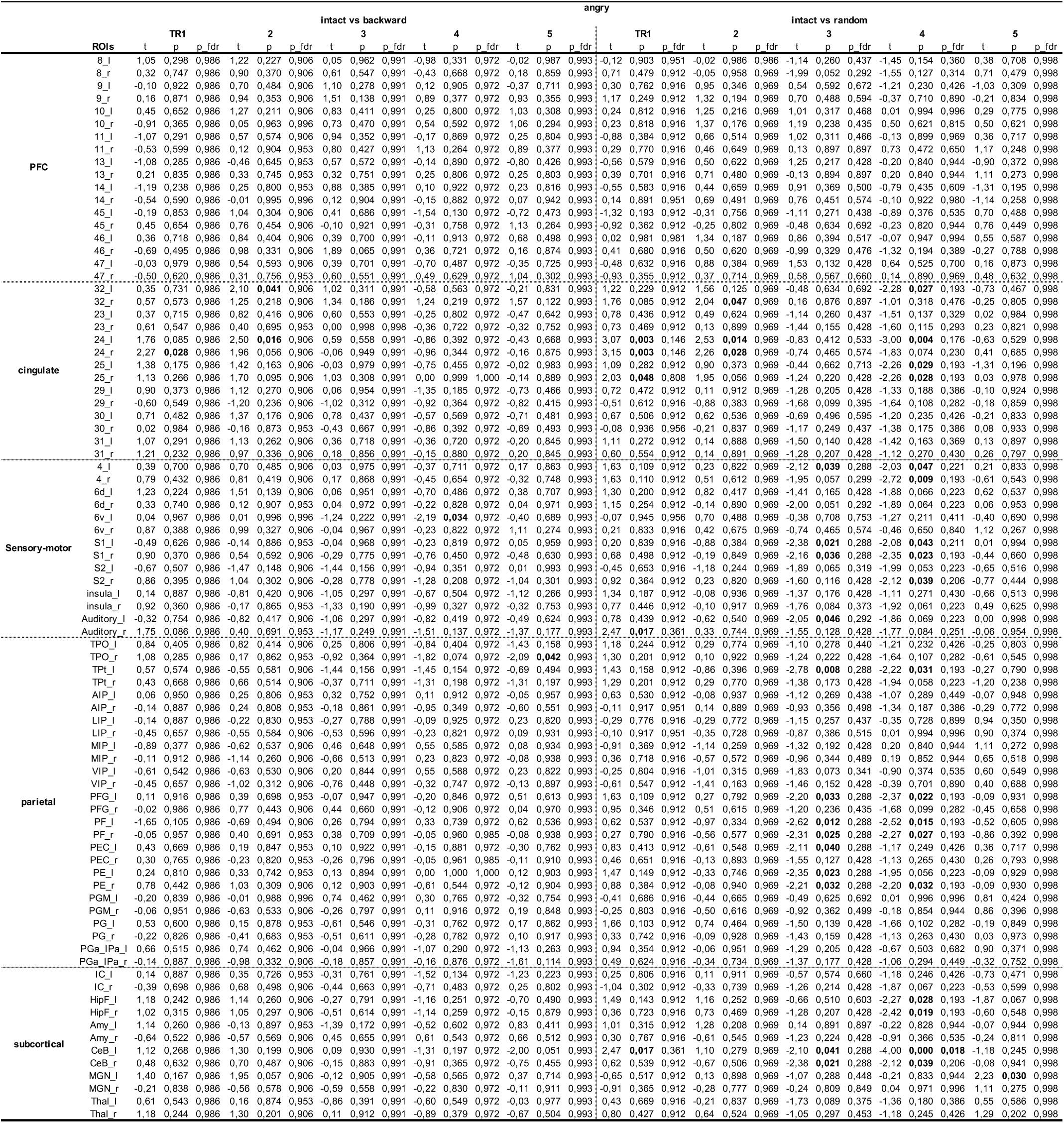

**supplementary Table 2.**
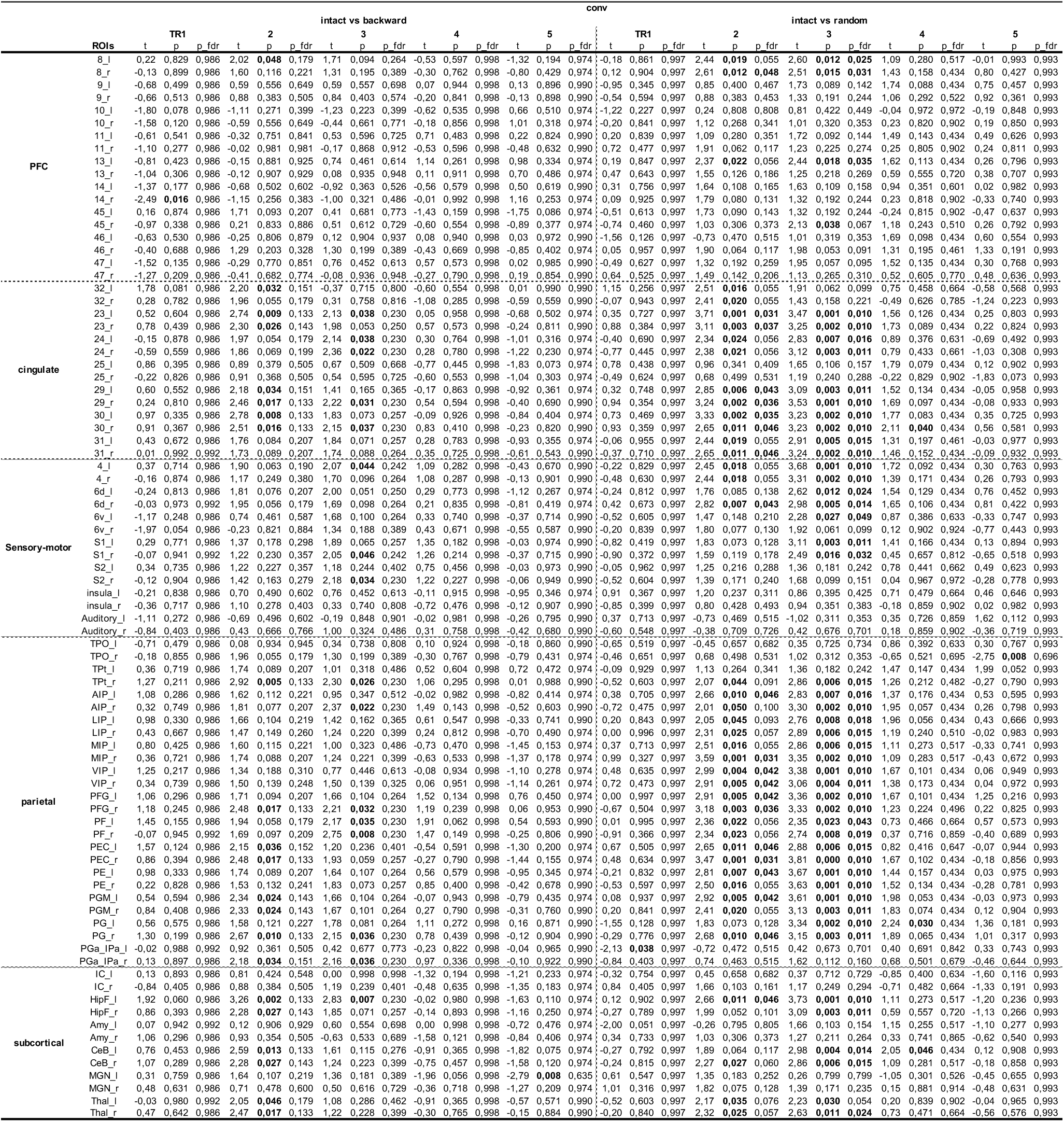

**supplementary Table 3.**
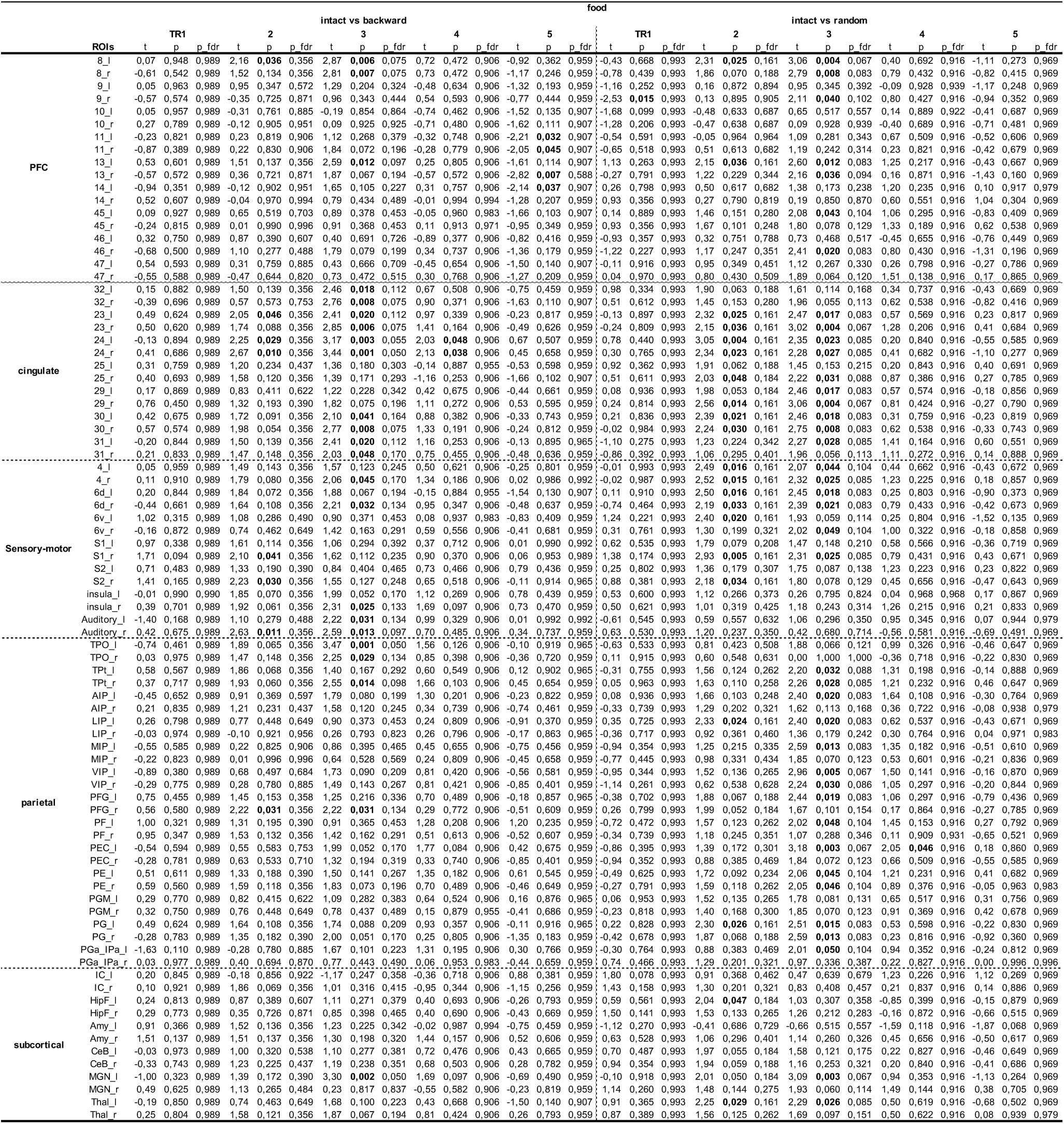

**supplementary Table 4.**

| percentile of significant ROIs |  | TR |  |  |  |  |
| --- | --- | --- | --- | --- | --- | --- |
|  |  | 1 | 2 | 3 | 4 | 5 |
| angry | intact vs backward | 1,19 | 2,38 | 0 | 1,19 | 1,19 |
|  | intact vs random | 5,95 | 3,57 | 15,48 | 21,43 | 1,19 |
| conv | intact vs backward | 1,19 | 26,19 | 19,05 | 0 | 1,19 |
|  | intact vs random | 1,19 | 50 | 57,14 | 3,57 | 1,19 |
| food | intact vs backward | 0 | 9,52 | 27,38 | 2,38 | 4,76 |
|  | intact vs random | 1,19 | 25 | 47,62 | 1,19 | 0 |

| Difference between TRs |  | angry |  |  |  |  |  | conv |  |  |  |  |  | food |  |  |  |  |  |
| --- | --- | --- | --- | --- | --- | --- | --- | --- | --- | --- | --- | --- | --- | --- | --- | --- | --- | --- | --- |
| TR | intact vs backward |  |  | intact vs random |  |  |  | intact vs backward |  |  | intact vs random |  |  | intact vs backward |  |  | intact vs random |  |  |
| | $\chi^2$ | p | p_fdr | $\chi^2$ | p | p_fdr | | $\chi^2$ | p | p_fdr | $\chi^2$ | p | p_fdr | $\chi^2$ | p | p_fdr | $\chi^2$ | p | p_fdr |
| 1 vs 2 | 0 | 1 | 1 | 0,25 | 0,617 | 0,617 |  | 17,39 | **<0.001** | **<0.001** | 37,21 | **<0.001** | **<0.001** | 6,13 | **0,013** | **0,027** | 16,41 | **<0.001** | **<0.001** |
| 1 vs 3 | 0 | 1 | 1 | 3,06 | 0,080 | 0,134 |  | 11,53 | **0,001** | **0,001** | 43,18 | **<0.001** | **<0.001** | 21,04 | **<0.001** | **<0.001** | 37,03 | **<0.001** | **<0.001** |
| 1 vs 4 | 0,5 | 0,48 | 1 | 8,47 | **0,004** | **0,009** |  | 0 | 1 | 1 | 0,25 | 0,617 | 0,617 | 0,5 | 0,480 | 0,533 | 0,5 | 0,480 | 0,599 |
| 1 vs 5 | 0,5 | 0,48 | 1 | 1,5 | 0,221 | 0,285 |  | 0,5 | 0,480 | 0,599 | 0,5 | 0,480 | 0,599 | 2,25 | 0,134 | 0,191 | 0 | 1 | 1 |
| 2 vs 3 | 0,5 | 0,48 | 1 | 5,06 | **0,024** | **0,049** |  | 1,14 | 0,286 | 0,409 | 2,5 | 0,114 | 0,163 | 10,32 | **0,001** | **0,003** | 12,96 | **<0.001** | **<0.001** |
| 2 vs 4 | 0 | 1 | 1 | 10,32 | **0,001** | **0,007** |  | 20,05 | **<0.001** | **<0.001** | 33,58 | **<0.001** | **<0.001** | 4,17 | **0,041** | 0,069 | 16,41 | **<0.001** | **<0.001** |
| 2 vs 5 | 0 | 1 | 1 | 0,25 | 0,617 | 0,617 |  | 17,39 | **<0.001** | **<0.001** | 37,21 | **<0.001** | **<0.001** | 0,75 | 0,386 | 0,483 | 19,05 | **<0.001** | **<0.001** |
| 3 vs 4 | 0 | 1 | 1 | 1,45 | 0,228 | 0,285 |  | 14,06 | **<0.001** | **<0.001** | 43,02 | **<0.001** | **<0.001** | 19,05 | **<0.001** | **<0.001** | 37,03 | **<0.001** | **<0.001** |
| 3 vs 5 | 0 | 1 | 1 | 8,64 | **0,003** | **0,009** |  | 11,53 | **0,001** | **0,001** | 43,18 | **<0.001** | **<0.001** | 12 | **0,001** | **0,002** | 38,03 | **<0.001** | **<0.001** |
| 4 vs 5 | 0,5 | 0,48 | 1 | 13,47 | **<0.001** | **0,002** |  | 0 | 1 | 1 | 0,25 | 0,617 | 0,617 | 0,17 | 0,683 | 0,683 | 0 | 1 | 1 |

| (intact vs backward) vs (intact vs random) |  | angry |  |  | conv |  |  | food |  |  |
| --- | --- | --- | --- | --- | --- | --- | --- | --- | --- | --- |
| TR | | $\chi^2$ | p | p_fdr | $\chi^2$ | p | p_fdr | $\chi^2$ | p | p_fdr |
| TR 1 |  | 2,25 | 0,134 | 0,223 | 0,5 | 0,480 | 0,480 | 0 | 1 | 1 |
| TR 2 |  | 0 | 1 | 1 | 13,88 | **<0.001** | **<0.001** | 8,47 | **0,004** | **0,017** |
| TR 3 |  | 11,08 | **0,001** | **0,002** | 26,69 | **<0.001** | **<0.001** | 7,31 | **0,007** | **0,017** |
| TR 4 |  | 13,47 | **<0.001** | **0,001** | 1,33 | 0,248 | 0,414 | 0 | 1 | 1 |
| TR 5 |  | 0,5 | 0,480 | 0,599 | 0,5 | 0,480 | 0,480 | 2,25 | 0,134 | 0,223 |

**supplementary Table 5.**
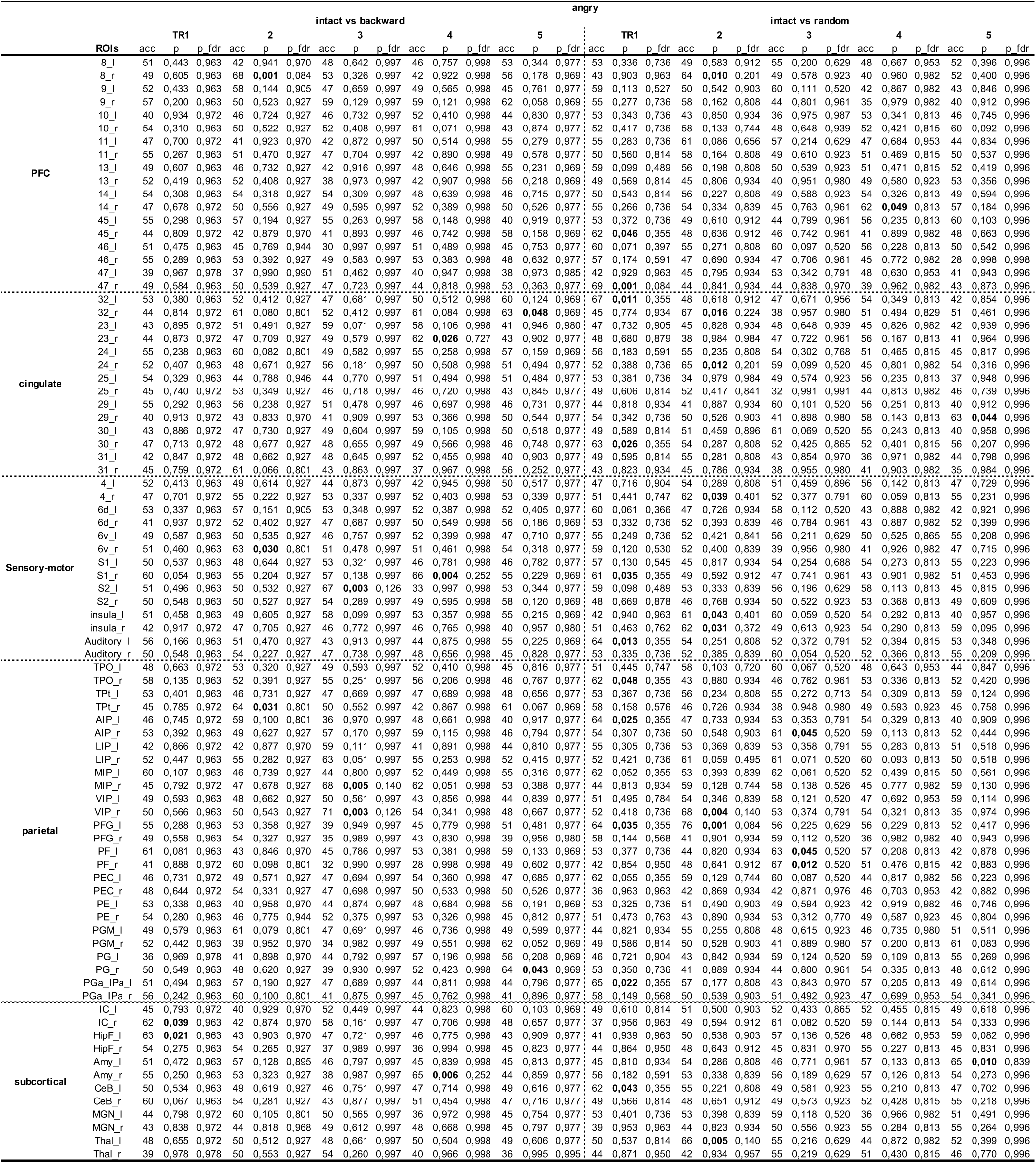

**supplementary Table 6.**
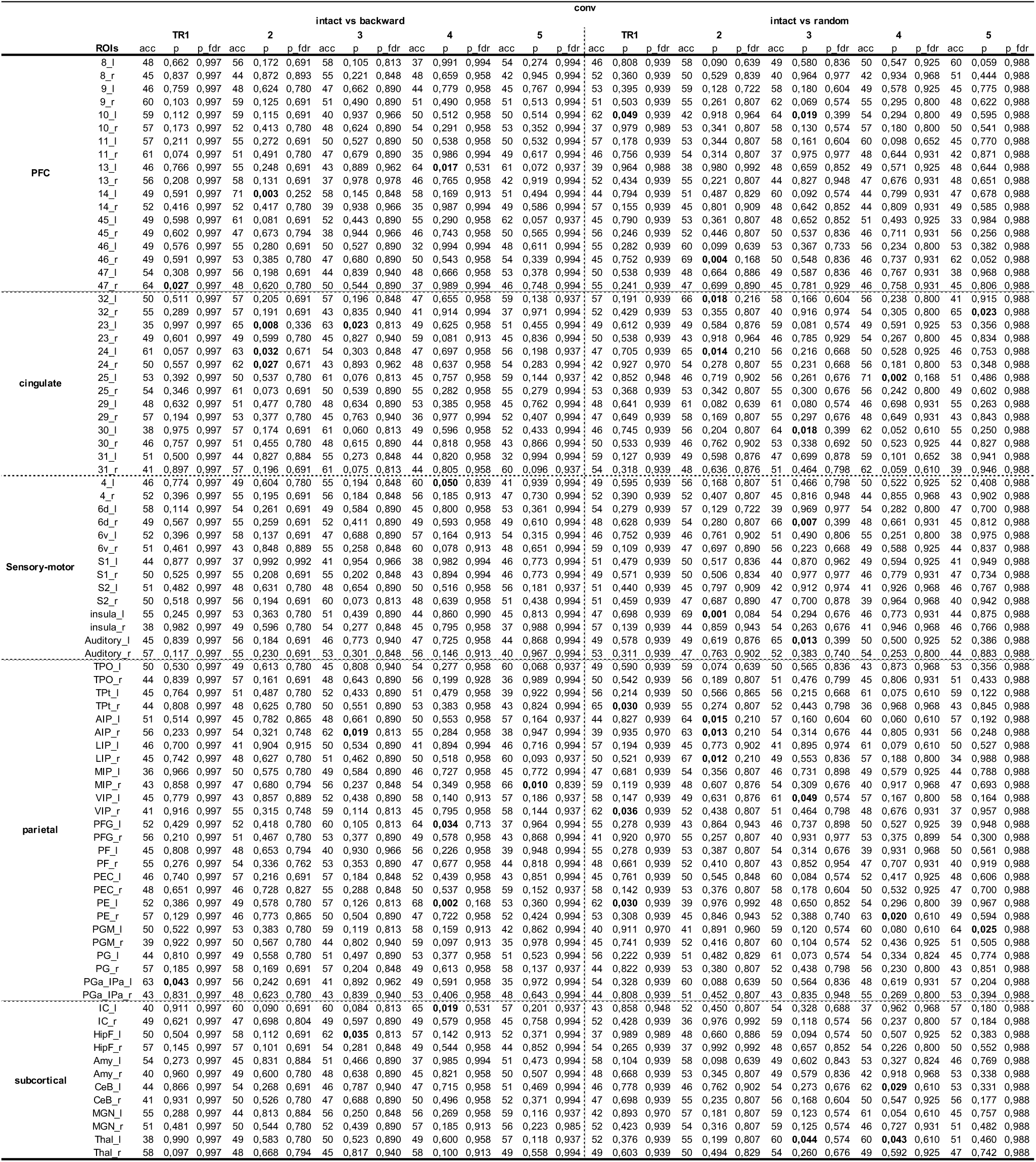

**Supplementary Table 7.**
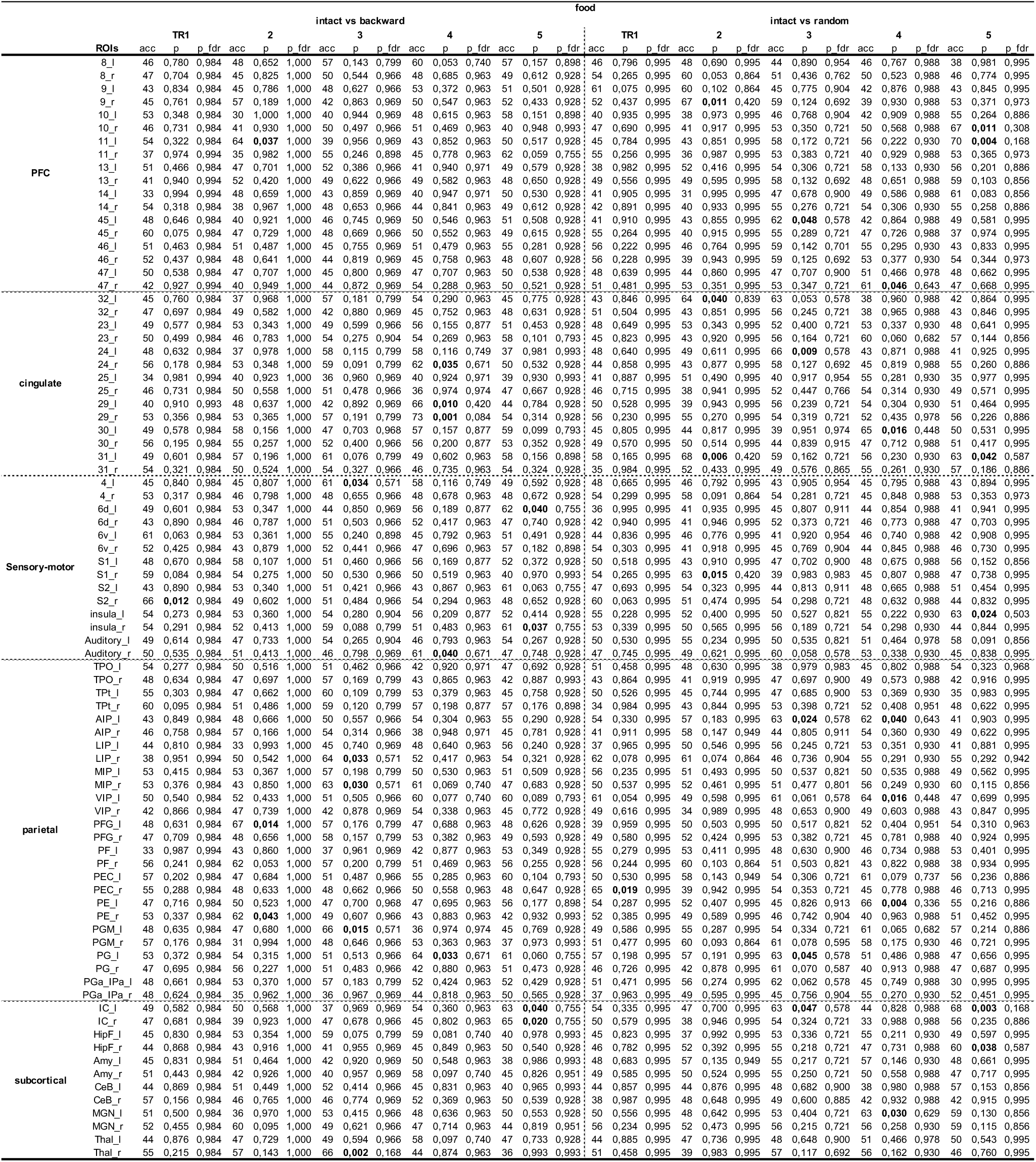

## Reference

1. Hasson, U., Yang, E., Vallines, I., Heeger, D.J., and Rubin, N. (2008). A hierarchy of temporal receptive windows in human cortex. J Neurosci 28, 2539–2550. 10.1523/JNEUROSCI.5487-07.2008.

2. Lerner, Y., Honey, C.J., Silbert, L.J., and Hasson, U. (2011). Topographic mapping of a hierarchy of temporal receptive windows using a narrated story. J Neurosci 31, 2906–2915. 10.1523/JNEUROSCI.3684-10.2011.

3. Honey, C.J., Thesen, T., Donner, T.H., Silbert, L.J., Carlson, C.E., Devinsky, O., Doyle, W.K., Rubin, N., Heeger, D.J., and Hasson, U. (2012). Slow cortical dynamics and the accumulation of information over long timescales. Neuron 76, 423–434. 10.1016/j.neuron.2012.08.011.

4. Simony, E., Honey, C.J., Chen, J., Lositsky, O., Yeshurun, Y., Wiesel, A., and Hasson, U. (2016). Dynamic reconfiguration of the default mode network during narrative comprehension. Nat Commun 7, 12141. 10.1038/ncomms12141.

5. Lahnakoski, J.M., Jääskeläinen, I.P., Sams, M., and Nummenmaa, L. (2017). Neural mechanisms for integrating consecutive and interleaved natural events. Hum Brain Mapp 38, 3360–3376. 10.1002/hbm.23591.

6. Baldassano, C., Hasson, U., and Norman, K.A. (2018). Representation of Real-World Event Schemas during Narrative Perception. J. Neurosci. 38, 9689–9699. 10.1523/JNEUROSCI.0251-18.2018.

7. Sabra, Z., Alawieh, A., Bonilha, L., Naselaris, T., and AuYong, N. (2022). Modulation of Spectral Representation and Connectivity Patterns in Response to Visual Narrative in the Human Brain. Front Hum Neurosci 16, 886938. 10.3389/fnhum.2022.886938.

8. Grall, C., Equita, J., and Finn, E.S. (2023). Neural unscrambling of temporal information during a nonlinear narrative. Cereb Cortex 33, 7001–7014. 10.1093/cercor/bhad015.

9. Assouline, A., and Mendelsohn, A. (2023). Weaving a story: Narrative formation over prolonged time scales engages social cognition and frontoparietal networks. European Journal of Neuroscience 57, 809–823. 10.1111/ejn.15909.

10. Zuo, X., Honey, C.J., Barense, M.D., Crombie, D., Norman, K.A., Hasson, U., and Chen, J. (2020). Temporal integration of narrative information in a hippocampal amnesic patient. Neuroimage 213, 116658. 10.1016/j.neuroimage.2020.116658.

11. Song, H., Park, B.-Y., Park, H., and Shim, W.M. (2021). Cognitive and Neural State Dynamics of Narrative Comprehension. J Neurosci 41, 8972–8990. 10.1523/JNEUROSCI.0037-21.2021.

12. Türker, B., Belloli, L., Owen, A.M., Naci, L., and Sitt, J.D. (2023). Processing of the same narrative stimuli elicits common functional connectivity dynamics between individuals. Sci Rep 13, 21260. 10.1038/s41598-023-48656-7.

13. Kolisnyk, M., Novi, S., Abdalmalak, A., Ardakani, R.M., Kazazian, K., Laforge, G., Debicki, D.B., and Owen, A.M. (2024). Assessing the consistency and sensitivity of the neural correlates of narrative stimuli using functional near-infrared spectroscopy. Imaging Neurosci (Camb) 2, imag–2–00331. 10.1162/imag_a_00331.

14. Zuberbühler, K., Cheney, D.L., and Seyfarth, R.M. (1999). Conceptual semantics in a nonhuman primate. Journal of Comparative Psychology 113, 33–42. 10.1037/0735-7036.113.1.33.

15. Arnold, K., and Zuberbühler, K. (2008). Meaningful call combinations in a non-human primate. Current Biology 18, R202–R203. 10.1016/j.cub.2008.01.040.

16. Ouattara, K., Lemasson, A., and Zuberbühler, K. (2009). Campbell’s monkeys concatenate vocalizations into context-specific call sequences. Proceedings of the National Academy of Sciences 106, 22026–22031. 10.1073/pnas.0908118106.

17. Miller, C.T., and Wang, X. (2006). Sensory-motor interactions modulate a primate vocal behavior: antiphonal calling in common marmosets. J Comp Physiol A 192, 27–38. 10.1007/s00359-005-0043-z.

18. Miller, C.T., Beck, K., Meade, B., and Wang, X. (2009). Antiphonal call timing in marmosets is behaviorally significant: interactive playback experiments. J Comp Physiol A 195, 783–789. 10.1007/s00359-009-0456-1.

19. Roy, S., Miller, C.T., Gottsch, D., and Wang, X. (2011). Vocal control by the common marmoset in the presence of interfering noise. J Exp Biol 214, 3619–3629. 10.1242/jeb.056101.

20. Takahashi, D.Y., Narayanan, D.Z., and Ghazanfar, A.A. (2013). Coupled oscillator dynamics of vocal turn-taking in monkeys. Curr Biol 23, 2162–2168. 10.1016/j.cub.2013.09.005.

21. Huang, J., Ma, H., Sun, Y., Chang, L., and Gong, N. (2022). Complex rules of vocal sequencing in marmoset monkeys. Preprint at bioRxiv, 10.1101/2022.08.03.502601 https://doi.org/10.1101/2022.08.03.502601.

22. Bosshard, A.B., Burkart, J.M., Merlo, P., Cathcart, C., Townsend, S.W., and Bickel, B. (2024). Beyond bigrams: call sequencing in the common marmoset (Callithrix jacchus) vocal system. R Soc Open Sci 11, 240218. 10.1098/rsos.240218.

23. Uhrig, L., Dehaene, S., and Jarraya, B. (2014). A Hierarchy of Responses to Auditory Regularities in the Macaque Brain. J. Neurosci. 34, 1127–1132. 10.1523/JNEUROSCI.3165-13.2014.

24. Wilson, B., Kikuchi, Y., Sun, L., Hunter, D., Dick, F., Smith, K., Thiele, A., Griffiths, T.D., Marslen-Wilson, W.D., and Petkov, C.I. (2015). Auditory sequence processing reveals evolutionarily conserved regions of frontal cortex in macaques and humans. Nat Commun 6, 8901. 10.1038/ncomms9901.

25. Kikuchi, Y., Attaheri, A., Wilson, B., Rhone, A.E., Nourski, K.V., Gander, P.E., Kovach, C.K., Kawasaki, H., Griffiths, T.D., Iii, M.A.H., et al. (2017). Sequence learning modulates neural responses and oscillatory coupling in human and monkey auditory cortex. PLOS Biology 15, e2000219. 10.1371/journal.pbio.2000219.

26. Nummela, S.U., Jovanovic, V., Mothe, L. de la, and Miller, C.T. (2017). Social Context-Dependent Activity in Marmoset Frontal Cortex Populations during Natural Conversations. J. Neurosci. 37, 7036–7047. 10.1523/JNEUROSCI.0702-17.2017.

27. Chao, Z.C., Takaura, K., Wang, L., Fujii, N., and Dehaene, S. (2018). Large-Scale Cortical Networks for Hierarchical Prediction and Prediction Error in the Primate Brain. Neuron 100, 1252–1266.e3. 10.1016/j.neuron.2018.10.004.

28. Jiang, Y., Komatsu, M., Chen, Y., Xie, R., Zhang, K., Xia, Y., Gui, P., Liang, Z., and Wang, L. (2022). Constructing the hierarchy of predictive auditory sequences in the marmoset brain. eLife 11, e74653. 10.7554/eLife.74653.

29. Grijseels, D.M., Fairbank, D.A., and Miller, C.T. (2024). A model of marmoset monkey vocal turn-taking. Proceedings of the Royal Society B: Biological Sciences 291, 20240150. 10.1098/rspb.2024.0150.

30. Park, J., Song, H., and Shim, W.M. (2025). Hippocampal systems for event encoding and sequencing during ongoing narrative comprehension. Commun Biol 8, 954. 10.1038/s42003-025-08377-1.

31. Naya, Y., and Suzuki, W.A. (2011). Integrating What and When Across the Primate Medial Temporal Lobe. Science 333, 773–776. 10.1126/science.1206773.

32. Regev, M., Honey, C.J., Simony, E., and Hasson, U. (2013). Selective and invariant neural responses to spoken and written narratives. J Neurosci 33, 15978–15988. 10.1523/JNEUROSCI.1580-13.2013.

33. Wang, F., and Diana, R.A. (2017). Neural correlates of temporal context retrieval for abstract scrambled phrases: Reducing narrative and familiarity-based strategies. Brain Res 1655, 128–137. 10.1016/j.brainres.2016.11.017.

34. Jafari, A., Dureux, A., Zanini, A., Menon, R.S., Gilbert, K.M., and Everling, S. (2023). A vocalization-processing network in marmosets. Cell Reports 42. 10.1016/j.celrep.2023.112526.

35. Dureux, A., Zanini, A., Trapeau, R., Belin, P., and Everling, S. (2025). Functional organization of voice patches in marmosets and cross-species comparisons with macaques and humans. Current Biology 35, 3869–3882.e4. 10.1016/j.cub.2025.07.008.

36. Nagarajan, G., Matrov, D., Pearson, A.C., Yen, C., Bradley, S.P., and Chudasama, Y. (2025). Cingulate cortex shapes early postnatal development of social vocalizations. eLife 13. 10.7554/eLife.97125.2.

37. Alexander, L., Wood, C.M., Gaskin, P.L.R., Sawiak, S.J., Fryer, T.D., Hong, Y.T., McIver, L., Clarke, H.F., and Roberts, A.C. (2020). Over-activation of primate subgenual cingulate cortex enhances the cardiovascular, behavioral and neural responses to threat. Nat Commun 11, 5386. 10.1038/s41467-020-19167-0.

38. Wilson, S.M., Saygin, A.P., Sereno, M.I., and Iacoboni, M. (2004). Listening to speech activates motor areas involved in speech production. Nat Neurosci 7, 701–702. 10.1038/nn1263.

39. Osnes, B., Hugdahl, K., and Specht, K. (2011). Effective connectivity analysis demonstrates involvement of premotor cortex during speech perception. Neuroimage 54, 2437–2445. 10.1016/j.neuroimage.2010.09.078.

40. Froesel, M., Ikuchi, K., Zhu, Q., Wang, H., Hauser, M., Ben Hamed, S., and Vanduffel, W. (2025). High-resolution fMRI reveals a dorsal brain pathway selective for conspecific vocalizations in macaques. Imaging Neuroscience 3, IMAG.a.108. 10.1162/IMAG.a.108.

41. Hage, S.R., and Nieder, A. (2016). Dual Neural Network Model for the Evolution of Speech and Language. Trends in Neurosciences 39, 813–829. 10.1016/j.tins.2016.10.006.

42. Zanini, A., Dureux, A., Jafari, A., Gilbert, K.M., Zeman, P., Bellyou, M., Li, A., Tuin, C.V., and Everling, S. (2023). *In vivo* functional brain mapping using ultra-high-field fMRI in awake common marmosets. STAR Protocols 4, 102586. 10.1016/j.xpro.2023.102586.

43. Gilbert, K.M., Dureux, A., Jafari, A., Zanini, A., Zeman, P., Menon, R.S., and Everling, S. (2023). A radiofrequency coil to facilitate task-based fMRI of awake marmosets. Journal of Neuroscience Methods 383, 109737. 10.1016/j.jneumeth.2022.109737.

44. Toarmino, C.R., Yen, C.C.C., Papoti, D., Bock, N.A., Leopold, D.A., Miller, C.T., and Silva, A.C. (2017). Functional magnetic resonance imaging of auditory cortical fields in awake marmosets. NeuroImage 162, 86–92. 10.1016/j.neuroimage.2017.08.052.

45. Cox, R.W. (1996). AFNI: software for analysis and visualization of functional magnetic resonance neuroimages. Comput Biomed Res 29, 162–173. 10.1006/cbmr.1996.0014.

46. Smith, S.M., Jenkinson, M., Woolrich, M.W., Beckmann, C.F., Behrens, T.E.J., Johansen-Berg, H., Bannister, P.R., De Luca, M., Drobnjak, I., Flitney, D.E., et al. (2004). Advances in functional and structural MR image analysis and implementation as FSL. Neuroimage 23 *Suppl 1*, S208–219. 10.1016/j.neuroimage.2004.07.051.

47. Liu, C., Ye, F.Q., Yen, C.C.-C., Newman, J.D., Glen, D., Leopold, D.A., and Silva, A.C. (2018). A digital 3D atlas of the marmoset brain based on multi-modal MRI. Neuroimage 169, 106–116. 10.1016/j.neuroimage.2017.12.004.

48. Glasser, M.F., Sotiropoulos, S.N., Wilson, J.A., Coalson, T.S., Fischl, B., Andersson, J.L., Xu, J., Jbabdi, S., Webster, M., Polimeni, J.R., et al. (2013). The minimal preprocessing pipelines for the Human Connectome Project. Neuroimage 80, 105–124. 10.1016/j.neuroimage.2013.04.127.

49. Coalson, T.S., Van Essen, D.C., and Glasser, M.F. (2018). The impact of traditional neuroimaging methods on the spatial localization of cortical areas. Proceedings of the National Academy of Sciences 115, E6356–E6365. 10.1073/pnas.1801582115.

50. Paxinos, G., Watson, C.R.R., Petrides, M., Rosa, M.G., and Tokuno, H. (2012). The Marmoset Brain in Stereotaxic Coordinates (Elsevier).

51. Hebart, M.N., Görgen, K., and Haynes, J.-D. (2015). The Decoding Toolbox (TDT): a versatile software package for multivariate analyses of functional imaging data. Front. Neuroinform. 8. 10.3389/fninf.2014.00088.

